# Matrix compliance regulates cellular and nuclear confinement of fibroblasts in tunable protein-based hydrogel microenvironments

**DOI:** 10.1101/2025.07.09.663931

**Authors:** Indira Priyadarshani Patra, Saujanya Sabarinath, Shiuly Sarkar, Abhijit Deshpande, Shantanu Pradhan

## Abstract

The extracellular matrix (ECM) influences cellular behavior, fate, and various mechanisms underlying homeostasis, development, and disease. Biophysical and biochemical properties of the ECM are known to affect three-dimensional (3D) cellular behavior, and phenotype that regulate a wide range of pathological conditions. Tunable biomimetic hydrogels are extensively employed to regulate cellular dynamics in defined 3D microenvironments, enabling detailed investigation of cell-matrix interactions. This study aims to establish the relationship between varying hydrogel properties (adhesivity, degradability, porosity, and stiffness, hereby collectively referred to as ‘matrix compliance’) and fibroblast morphology at the cellular and nuclear levels. Poly(ethylene glycol diacrylate)-fibrinogen (PF)-based hydrogels were fabricated and their matrix properties were tuned using two different non-degradable co-monomers of varying molecular weights and concentrations. NIH3T3 mouse fibroblasts were cultured in 3D hydrogels for 10 days and their cellular and nuclear morphology was analyzed via confocal imaging. Cells in degradable, soft, porous, and adhesive hydrogels displayed higher degree of spreading, higher cell density, longer protrusions, and elongated and larger nuclei compared to those in less degradable, stiffer, less porous, and less adhesive hydrogels. Matrix compliance conjointly regulated cell density, protrusion length, and protrusion frequency (collectively referred to as ‘cellular confinement’) and nuclear volume (referred to as ‘nuclear confinement’). These results highlight the complex interplay and interdependence of these cellular and nuclear morphological features as regulated by engineered tunable matrices, providing guidance for future studies of 3D cell behavior in novel biomaterials.

**Statement of significance:** Biomimetic tunable hydrogels are commonly used to support, and control, three-dimensional (3D) cellular behavior to recapitulate various developmental processes and disease states. Specific biophysical and biochemical characteristics of the extracellular matrix (ECM) can be modeled using these hydrogels by controlling the compositions and crosslinking mechanisms. Using this approach, the behavior of cells in 3D hydrogel matrices can be correlated with the matrix properties, thereby providing mechanistic insights into cell-matrix interactions. This study assesses the variations in the morphological features of fibroblasts encapsulated in 3D protein-based hydrogel matrices with varying crosslinking mechanisms. Our results reveal the combinatorial role of matrix adhesivity, degradability, porosity, and stiffness in regulating cellular and nuclear confinement of fibroblasts in 3D microenvironments. Overall, this study elucidates how matrix characteristics influence cellular behavior, serving as a practical guide for researchers developing innovative biomaterial matrices for 3D cell culture applications.

## 1. INTRODUCTION

The extracellular matrix (ECM) is a major regulator of a wide variety of cellular processes associated with development, morphogenesis, cellular structure and function, pathological progression and ultimately treatment outcomes [1–3]. The role of the dysregulated ECM in the context of various pathologies (cancer, fibrosis, aging, inflammation etc.) has been widely studied [4,5]. With regard to the ECM, specific biochemical and biophysical properties (composition of the matrisome, stiffness, porosity, isotropy) have been correlated to cellular behavior and phenotypic outcomes. In previous studies, the role of biophysical features of the matrix have been used to study cell spreading, adhesion, migration, differentiation, and other morphological characteristics with model cell lines including fibroblasts, endothelial cells, smooth muscle cells, stem cells and others [6–10]. Although studies of two-dimensional (2D) cell-seeded substrates are relatively straightforward, studies in three-dimensional (3D) matrices with encapsulated cells involve several intertwined biological variables, decoupling of which is very challenging. Particularly, specific contributions of each of these matrix parameters on cell fate and state is yet to be elucidated.

In general, encapsulated cells within 3D matrices need to overcome one of four rate-limiting steps towards cellular morphogenesis: 1) the ability to adhere/attach to the matrix ligands, 2) the ability to proteolytically degrade the matrix to create niches for filopodial protrusions, 3) the ability to undergo plastic deformation through the available porous network, and 4) the ability to generate sufficient traction force on the matrix to overcome yield stress and physically deform the matrix [11,12]. The ability of cells to adhere to the matrix depends on the density and accessibility of adhesive ligands in the backbone of the matrix-constituent and the density of integrins with optimal conformation/clustering on the cell surface. The ability of cells to proteolytically degrade the matrix depends on the density and accessibility of proteolytically cleavable sites in the network backbone and the relative expression and secretion of matrix metalloproteases (MMPs). In the absence of integrin-mediated cell adhesion or poorly degradable matrix, cells can employ traction force-mediated physical stretching or pushing of the matrix network to switch from a mesenchymal mode to amoeboid mode of migration [13,14]. The amoeboid mode of migration and spreading is further dictated by the matrix porosity and matrix stiffness (defined by yield strength of the network against cell-generated traction forces). Hence, to effectively correlate matrix structure with 3D cellular response and behavior, it is essential to characterize, at a minimum, four key matrix features: adhesivity, porosity, degradability, and stiffness.

To investigate and establish the role of ECM features on 3D cell behavior, hydrogels have emerged as a versatile biomaterial of choice due to its amenability for cell culture, robust control of engineered features to simulate native microenvironmental conditions, and ease of handling, observation, and analysis [15]. In past studies, the most abundantly occurring natural ECM component, collagen I, has been tuned with respect to matrix density, fiber alignment, thickness and length to quantify 2D and 3D cell behavior [16,17]. Other natural ECM-mimetic materials used for 2D and 3D cellular investigations include protein-based (fibrin, gelatin, silk etc.) or polysaccharide-based (alginate, hyaluronic acid etc.). hydrogels, while some synthetic biomaterials include poly(ethylene glycol)-based derivatives, poly(lactic-co-glycolic) acid, amongst others [18–20]. In this study, a biosynthetic matrix was used comprising of fibrinogen (natural ECM component) conjugated with poly(ethylene glycol diacrylate) (PEGDA) to obtain PEG-fibrinogen (PF) (Figure 1A) [21,22]. Fibrinogen provides the cell-adhesive and proteolytically degradable sites required for time-dependent 3D cellular remodeling, while PEGDA provides stability, structural integrity, and mechanical support to the overall matrix through crosslinking of the protein chains. The PEG-protein network can be further strengthened by incorporating functional co-monomers that can introduce additional crosslinks (including the formation of double-networks) within the existing network through free-radical polymerization [23].

**Figure 1:**
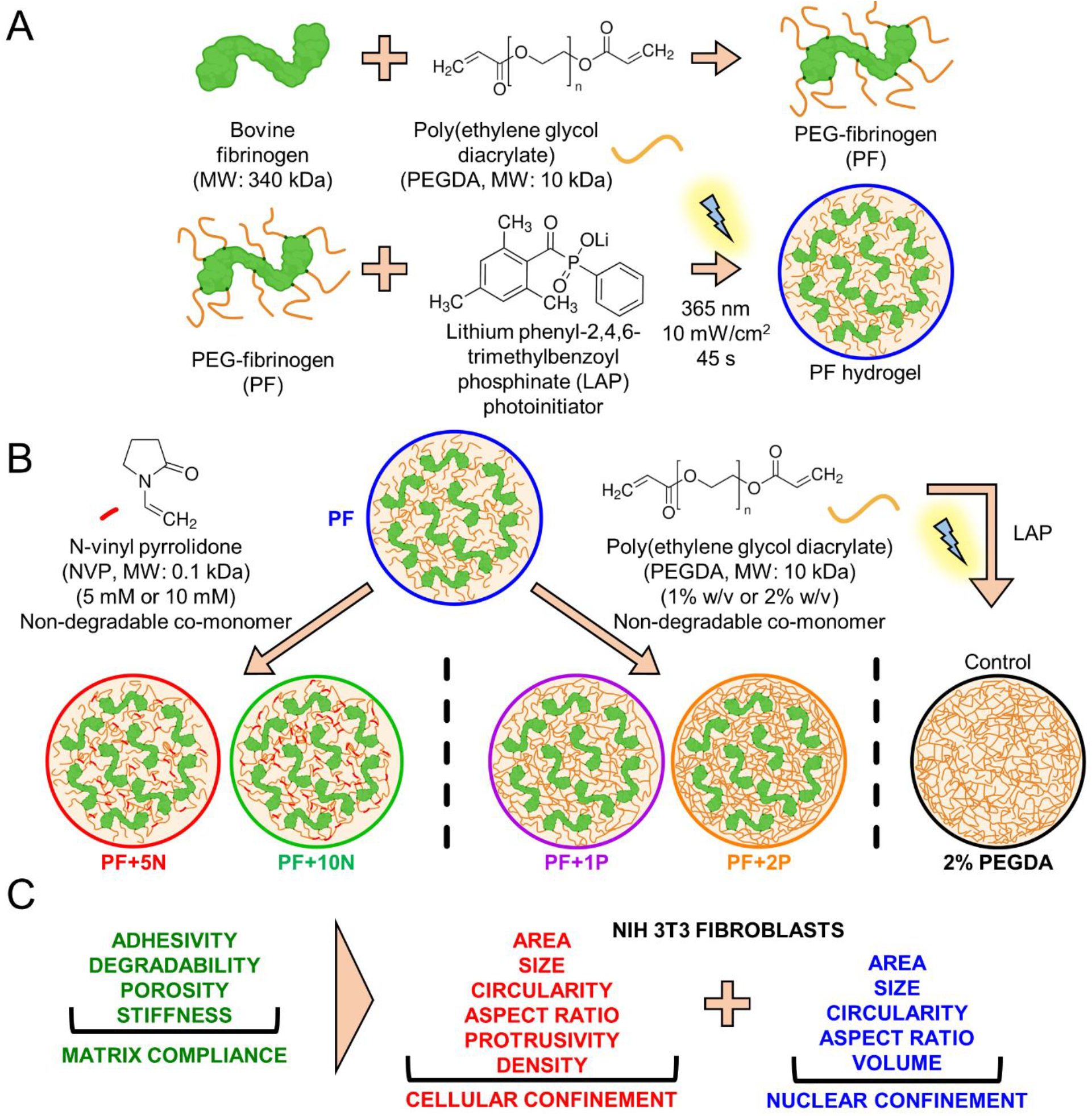
Schematic of experimental workflow. (A) Fibrinogen is covalently conjugated with PEGDA to synthesize PEG-fibrinogen (PF). PF precursor is photocrosslinked with LAP under light (365 nm) to form PF hydrogel. (B) PF precursor is mixed with additional non-degradable low molecular weight co-monomer NVP (5 or 10 mM) or non-degradable co-monomer high molecular weight PEGDA (1% or 2% w/v) to form hydrogels of varying compositions. As control, PEGDA precursor is also crosslinked to form PEGDA hydrogel. Created in BioRender. Pradhan, S. (2025) (C) Hydrogels are characterized to determine matrix adhesivity, degradability, porosity, and stiffness (collectively termed as ‘matrix compliance’). NIH3T3 mouse fibroblasts are cultured in 3D hydrogels and various morphological parameters are evaluated to correlate cellular and nuclear confinement with matrix compliance.

Specifically, the existing PF network has been modified by introducing N-vinyl-pyrrolidone (NVP) (111 Da) or PEGDA (10 kDa) as non-degradable co-monomers that conjointly affect matrix biophysical and biochemical properties, with only PEGDA serving as the control matrix (Figure 1B). Four distinct hydrogel properties: adhesivity, degradability, porosity, and stiffness, were quantified for each of the six hydrogel compositions and collectively termed as ‘matrix compliance’. NIH3T3 fibroblasts were encapsulated in 3D hydrogels and their cellular and nuclear features were quantified as a function of the matrix properties (Figure 1C). PF and PF with low concentration of NVP were observed to be permissive, compliant matrices and allowed high degree of cell spreading and protrusivity (low cellular confinement) and nuclear elongation and volume (low nuclear confinement), while PF with high concentration of PEGDA and control PEGDA matrix were observed to be restrictive (poorly compliant) matrices with low cellularity and protrusivity (high cell confinement) and rounded, smaller nuclei (high nuclear confinement). These approaches and results provide a theoretical framework for understanding the role of matrix features in driving cellular (and subsequently tissue-scale) morphogenesis, which can be used to guide a wide variety of applications in designing novel biomaterials for tissue engineering, and for mechanistic investigations of cell-ECM interactions in cancer, fibrosis, and other diseases.

## 2. MATERIALS AND METHODS

### 2.1 Cell culture and maintenance

NIH3T3 mouse fibroblasts were obtained from National Centre for Cell Science, Pune. Cells were maintained in DMEM (Himedia), supplemented with 10% fetal bovine serum (GIBCO^®^, Thermo Fisher Scientific), 1% (v/v) L-Glutamine (Himedia), 1% (v/v) sodium pyruvate (Himedia), 1% (v/v) penicillin/streptomycin (Himedia). Cells from passage 7-8 were used for experiments.

### 2.2 Hydrogel synthesis and characterization

PEGDA (molecular weight: 10 kDa) was synthesized and characterized by ^1^H NMR spectroscopy as described previously [22,23]. The lyophilized PEGDA was covalently conjugated to bovine fibrinogen (Sigma-Aldrich) to form PEG-fibrinogen (PF) as per established protocol [22,23]. Details are provided in Supplementary Methods. The synthesized PF was characterized for the protein content (using BCA assay, Pierce™ BCA Protein Assay Kit, Thermo-Scientific) and the PEGDA content (lyophilization and weighing of fixed weight of product). The PEGylation efficiency was calculated according to the below formula as given below in Equation(1).

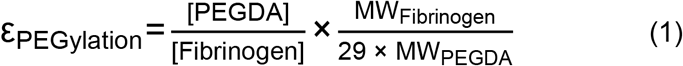

### 2.3 Hydrogel crosslinking and cell encapsulation

Uniform circular molds for hydrogel fabrication were prefabricated by curing a poly(dimethyl siloxane) (PDMS) sheet between two glass slides separated by 1 mm thick spacers, followed by punching with a 4 mm biopsy punch. To prepare the hydrogel precursor, PF (in phosphate buffered saline, PBS) solution was mixed with UV-sensitive photoinitiator, lithium phenyl 2,4,6-trimethylbenzoylphosphinate (LAP) (synthesized separately, see Supplementary Methods), at a final concentration of 10 mM. 1-vinyl-2-pyrrolidone (NVP, Sigma-Aldrich) was dissolved in PBS at 500 mM stock and mixed with the precursor to a final concentration of 5 mM or 10 mM to obtain PF+5N or PF+10N hydrogel compositions. PEGDA powder (10 kDa) was dissolved in PBS at 300 mg/mL stock and mixed with the precursor to a final concentration of 1% w/v or 2% w/v to obtain PF+1P and PF+2P hydrogel compositions. The concentrations of excess PEGDA added was adjusted based on the degree of acrylation estimated from ^1^H NMR. 10 μL of the polymer precursor solution was pipetted into each PDMS mold affixed in a well plate and photocrosslinked under UV light (Blak Ray B100 Spot UV Lamp, 365 nm, 10 mW/cm^2^) for 45 seconds to form fully crosslinked gels. After crosslinking, PDMS molds were gently peeled away, leaving behind disc-shaped crosslinked hydrogels, which were used for further assays. For cellular studies, NIH3T3 fibroblasts cultured in T25 flasks were trypsinized, counted, and resuspended in the polymer precursor at a concentration of 5×10^6^ cells/mL before crosslinking. Post crosslinking, the encapsulated cells in the free-floating hydrogels were maintained in DMEM media for ten days, with media changes every two days.

### 2.4 Swelling studies

Hydrogels of varying compositions (discs of 4 mm diameter, 1 mm thickness) were crosslinked as described above. Immediately after crosslinking, hydrogels were carefully weighed on a tared analytical balance and their pre-swollen weights were recorded. The hydrogels were then incubated in PBS for 6 hours to reach equilibrium swelling and reweighed to measure their swollen weight. Five replicates for each hydrogel composition were measured using Equation (2).

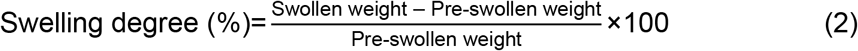

### 2.5 Hydrogel diffusion and porosity

The porosity (indicated by the theoretical mesh size) of the hydrogels of varying compositions was measured using macromolecular diffusion assay as reported earlier [23]. Briefly, photocrosslinked hydrogels were swollen to equilibrium and incubated overnight in tetramethylrhodamine-5-isothiocyanate (TRITC)-dextran (molecular weight 4.4 kDa or 150 kDa, Thermo Fisher Scientific) in PBS (0.5 mg/mL). The loaded hydrogels were transferred to PBS buffer and the release of TRITC-dextran from the loaded hydrogel into the surrounding buffer was measured at 15-minute intervals for 3 hours using a multi-mode plate reader (Biotek Synergy H1). A minimum of 5 hydrogel replicates per condition were measured. Details are provided in Supplementary Methods.

### 2.6 Hydrogel degradation

The relative degradability of hydrogels of varying composition was measured using enzymatic degradation under collagenase I over time as reported earlier [23]. Briefly, photocrosslinked hydrogels were swollen to equilibrium and incubated in Coomassie Brilliant Blue G-250 dye (Sigma-Aldrich) for protein-specific staining. The stained hydrogels were incubated in destaining buffer to remove any unbound dye. The destained hydrogels were incubated with collagenase I (Himedia) in PBS buffer at 37°C to initiate degradation. The enzymatically degraded Coomassie-bound protein was collected at 15-minute intervals for 3 hours and the absorbance (595 nm) was measured using a multimode plate reader (Biotek Synergy H1). A minimum of 5 hydrogel replicates per condition were measured. Details are provided in the Supplementary Methods.

### 2.7 Fourier transform infrared spectroscopy

In order to confirm the successful incorporation of the co-monomers and modification of the base PF hydrogel, FTIR spectroscopy was conducted on the hydrogel samples. Briefly, photocrosslinked hydrogels and precursors were lyophilized and prepared as pellets, which were measured against pure potassium bromide (KBr) pellets as a blank reference. FTIR spectra were recorded in transmittance mode across a wavenumber range of 4000 cm^-1^ to 400 cm^-1^. To ensure accurate analysis, the spectra for all samples were normalized, with major vibrational bands assigned to their respective chemical bonds, which facilitated the identification and confirmation of specific functional groups present in each sample.

### 2.8 Scanning electron micrography

Scanning electron micrography (SEM) of the crosslinked hydrogels was conducted to assess variations in the ultrastructural morphology. Briefly, the photocrosslinked hydrogels were swollen to equilibrium in PBS, cryo-frozen in liquid nitrogen for 30 seconds, and subsequently dried via lyophilization for 48 hours. Once dried, the samples were mounted on carbon taped-aluminum stubs, sputter-coated with gold and imaged using a scanning electron microscope (FEI-Quanta FEG 200F).

### 2.9 Rheological characterization

The mechanical properties of the hydrogels of varying compositions were characterized using rheometry. Briefly, photocrosslinked hydrogels were swollen to equilibrium and loaded on a rheometer (Anton Paar MCR302e) at 37°C, equipped with a Peltier plate temperature-controlled base and an 8 mm parallel plate configuration. To ensure uniform sample size, hollow punches with an 8 mm diameter were used to obtain consistent 3D disc-shaped hydrogels. These were carefully placed flat on the rheometer’s bottom plate (sample platform) for testing. To determine the linear viscoelastic range, amplitude sweep measurements were taken by varying the strain from 0.1% to 100%, while the frequency remained constant at 1 rad/s. For the frequency sweep measurements, the hydrogels were tested under a 1% strain, with an oscillation frequency (ω) ranging from 0.1 to 100 rad/s. Three hydrogel replicates for each condition were tested. The yield stress was determined from the intersection of the storage and loss modulus of the amplitude sweep measurements. The complex shear modulus was determined from the storage and loss moduli obtained from the frequency sweep given by equation (3).

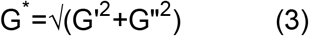

The complex viscosity (η*) was determined using equation (4).

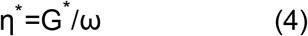

The loss tangent was calculated from the frequency sweep using equation (5).

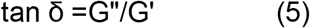

The compressive modulus (E) was estimated using equation (6).

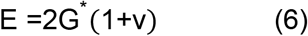

where v= Poisson’s ratio (assumed to be 0.5 for viscoelastic hydrogels)

Further, based on the swollen polymer network model, the shear modulus each of the hydrogel compositions was estimated and compared with the experimental values obtained by Equation (3). Details of the model equations and parameters are provided in Supplemental Methods.

### 2.10 Fluorescence staining and confocal imaging

The 3D morphology of encapsulated NIH3T3 fibroblasts was visualized through fluorescence staining and confocal fluorescence microscopy. Cell-laden hydrogels, maintained in 3D culture for 10 days, were carefully washed with PBS and fixed with 4% paraformaldehyde (Himedia) for 1 hour at 25°C. Permeabilization was carried out using 0.5% Triton-X for 30 minutes, followed by incubation in blocking buffer (2% bovine serum albumin and 5% fetal bovine serum in PBS) for 30 minutes. Hydrogels were incubated with Alexa Fluor^®^ 568 Phalloidin (Invitrogen, 25 μL/mL in blocking buffer) for 1 hour to stain F-actin, followed by nuclear staining with Hoechst 33342 (10 μL/mL in blocking buffer) for 30 minutes. Hydrogels were washed with PBS and transferred to glass-bottom dishes for confocal imaging (Olympus Fluoview 3000). Z-stacks of 200 μm thickness were acquired, with individual slices spaced 5 μm apart. A minimum of three independent z-stacks from three independent hydrogels were imaged per condition.

### 2.11 Image analysis

Cellular and nuclear morphology was analyzed from fluorescent confocal z-stacks and processed in FIJI software (NIH, Version 1.54p). Cell density was quantified by thresholding and automated nuclei counting in a defined volume of the z-stack. F-actin positive cells were quantified by manual counting of cells within the z-stack. The protrusion length and protrusion frequency were measured by manually drawing lines along the F-actin-stained protrusions from the cell bodies. In each z-stack, individual cells were outlined manually, and their morphological features (area, perimeter, major and minor axis, circularity and aspect ratio) were measured. Similarly, for nuclear quantification, the z-stacks were thresholded and the nuclear features were extracted in an automated manner. A minimum of 200 cells or nuclei were quantified for each condition. The mean geometric diameter of cell (or nuclei) was estimated by the equation below.

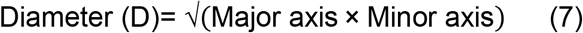

The effective nuclear volume was quantified as [24]:

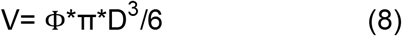

where Φ is the nuclear sphericity defined as:

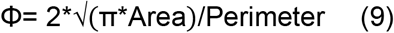

The matrix adhesivity (normalized fibrinogen content), degradability (normalized degradation rate), porosity (normalized mesh size), and stiffness (normalized compressive modulus) were used to calculate a unified metric, matrix compliance (ranging from 0.0 to 1.0). The normalized cell density, normalized protrusion length, and normalized protrusion frequency were used to calculate ‘cell confinement’, while the normalized effective nuclear volume was used to calculate nuclear confinement.

### 2.12 Statistical analysis

Statistical analysis was performed using GraphPad Prism 9.5.1 software. For multiple comparison across groups, normality of the Gaussian distribution was verified prior to one-way ANOVA test, with a Tukey’s family error rate of 5% was employed to assess statistical significance between different groups, assuming equal variance and sample size of the compared groups. Unless otherwise specified, p<0.05 was regarded as statistically significant.

## 3. RESULTS

### 3.1 Hydrogel composition

Fibrinogen-based biosynthetic hydrogel matrices of varying compositions were formed to investigate the relationship between cellular and nuclear features as a function of hydrogel characteristics. PEG-fibrinogen (PF) hydrogels were synthesized by covalent conjugation of poly(ethylene glycol diacrylate) (PEGDA) and fibrinogen (Fb) via Michael’s addition reaction. The degree of acrylation of PEGDA, quantified through NMR, was estimated to be 73% and the degree of PEGylation of Fb was estimated to be 138%. The higher degree of PEGylation can either be attributed to reaction of PEGDA with additional amino acid residues other than cysteine or to small errors in concentration estimations of protein or PEGDA. Nevertheless, PF hydrogels had a protein content of 12.0 mg/mL and PEGDA content of 14.2 mg/mL at a molar ratio of 40:1 (PEGDA:Fb) (Table 1). The available number of moles of free acrylate (those not bound to the Fb cysteines) was calculated to be 0.4×10^-7^/mL, which gives an indirect estimate of the crosslinkability of the hydrogel matrix.

**Table 1:** Estimated values of hydrogel compositions.

| Parameter/<br>Composition | PF | PF+5N | PF+10N | PF+1P | PF+2P | PEGDA |
| --- | --- | --- | --- | --- | --- | --- |
| Fibrinogen concentration (mg/mL) | 12.0 | 12.0 | 12.0 | 11.6 | 11.2 | 0.0 |
| PEGDA concentration (mg/mL) | 14.2 | 14.2 | 14.2 | 23.7 | 33.2 | 20.0 |
| NVP concentration (mM) (μL/mL) | 0.0<br>(0.0) | 5.0<br>(0.5) | 10.0<br>(1.0) | 0.0<br>(0.0) | 0.0<br>(0.0) | 0.0<br>(0.0) |
| Total mass of polymer (%w/v) | 2.6 | 2.6 | 2.6 | 3.5 | 4.4 | 2.0 |
| Molar ratio of PEGDA: fibrinogen | 40:1 | 40:1 | 40:1 | 70:1 | 101:1 | - |
| Molar ratio of NVP: free acrylate | - | 7:1 | 14:1 | - | - | - |
| Crosslinking density (moles of free acrylate/mL) | $0.4 \times 10^{-7}$ | $0.4 \times 10^{-7}$ | $0.4 \times 10^{-7}$ | $2.3 \times 10^{-7}$ | $5.0 \times 10^{-7}$ | $4.0 \times 10^{-7}$ |

With addition of low concentration of 5 mM or 10 mM N-vinyl-pyrrolidone (NVP), the protein and PEGDA content (and thereby the total polymer content) and the available moles of free acrylate remained constant, due to the low molecular weight of NVP (111 g/mol). However, due to the presence of the vinyl group in NVP (a non-degradable co-monomer), the junction functionality of the polymer chains was increased and the free acrylates of PEGDA were able to form stronger and less-degradable crosslinks with each other. In comparison, addition of 1% w/v or 2% w/v of 10 kDa non-degradable PEGDA to the PF network led to significant increase in the total PEGDA content (with a slight decrease in the protein content), significant increase in total polymer content and hence the available moles of free acrylate for crosslinking. The additional PEGDA formed crosslinks both with the existing free acrylates of the Fb-conjugated PEGDA and with each other, thereby forming a double interpenetrating network within the hydrogel matrix. As control, PEGDA hydrogels (2% w/v), without any protein content were also formed to obtain a hydrogel matrix with comparable polymer content and crosslinking density.

### 3.2 Hydrogel swelling

To test the effect of additional crosslinking within PF hydrogel networks, fabricated hydrogels were tested for swellability. At equilibrium swelling, PF hydrogels swelled to 81±7.2% of its initial weight post crosslinking (Figure 2A). With additional NVP, the swelling degree was reduced to 44±3.5% and 43±2.0% for PF+5N and PF+10N hydrogels respectively. This was primarily due to additional crosslinks of the PEGDA chains with the vinyl groups of NVP, which significantly reduced the chain flexibility of the crosslinked polymer network, resulting in reduced capacity to absorb water in its equilibrium swollen state. In contrast, additional PEGDA increased the swelling degree to 100±2.3% and 113±3.1% for PF+1P and PF+2P hydrogels respectively. As control, PEGDA hydrogels (without any protein) displayed a swelling degree of 120±1.8%. These results demonstrate that incorporation of hydrophilic PEGDA in the protein-polymer crosslinked network significantly increased junction functionality and polymer chain flexibility to retain water in the equilibrium swollen state. For the same chemical composition, the degree of swelling is inversely related to the crosslink density. Therefore, these results indicate that effective crosslink density of these hydrogels is not correlated with the free acrylate due to differing interactions in different compositions.

**Figure 2:**
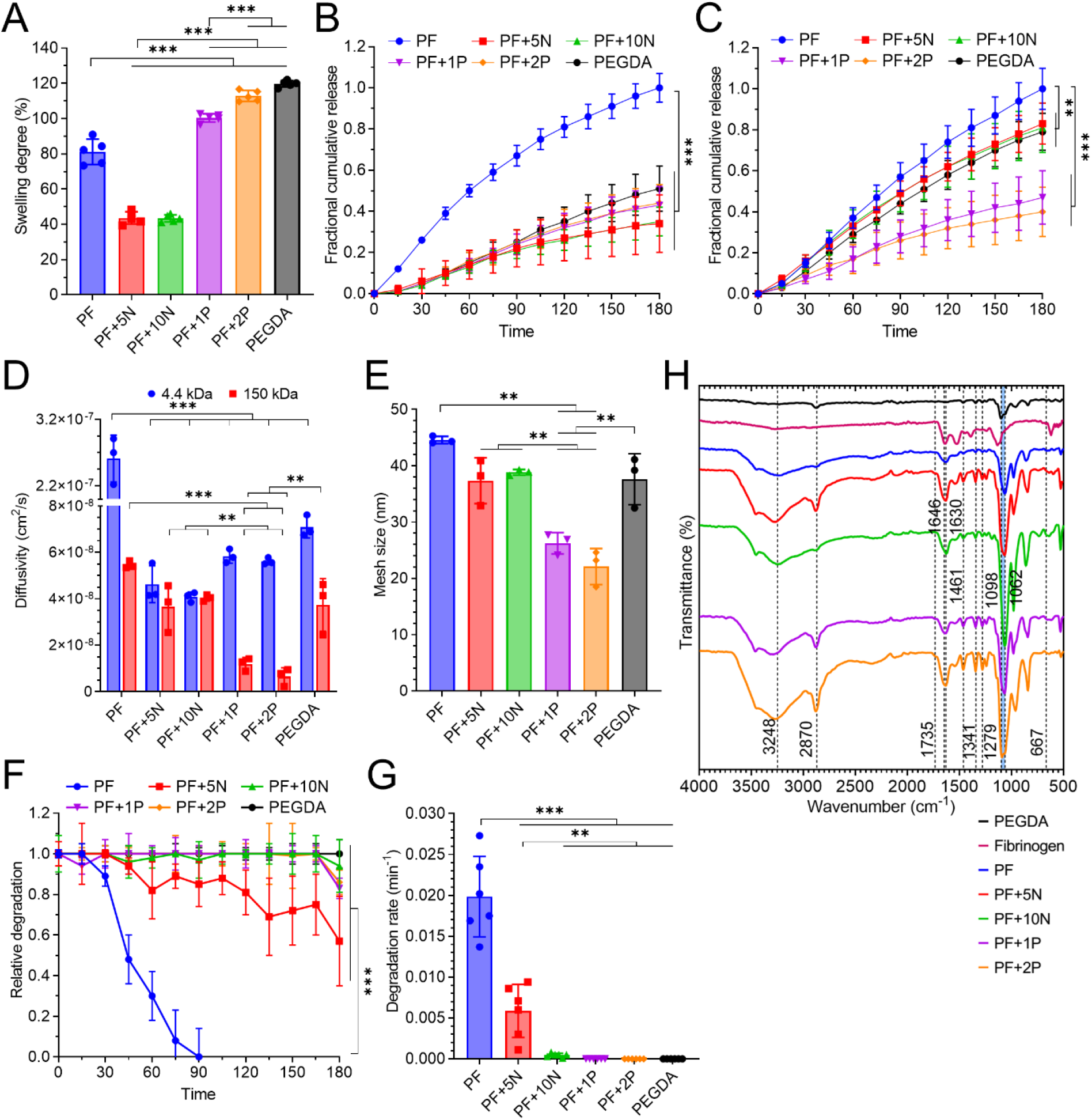
Hydrogel characterization. (A) Degree of equilibrium swelling of hydrogels. PF hydrogels with additional NVP had reduced swelling while those with additional PEGDA displayed higher swelling compared to PF hydrogels. (B) Relative cumulative release of TRITC-dextran (MW: 4.4 kDa) and (C) TRITC-dextran (MW: 150 kDa) for various hydrogel compositions. (D) Estimated diffusivities of 4.4 kDa and 150 kDa TRITC-dextran. PF hydrogels with additional PEGDA have reduced diffusivity of 150 kDa TRITC-dextran. (E) Calculated theoretical mesh size of hydrogels from diffusivity values. Hydrogels become less porous with addition of excess PEGDA. (F) Relative degradation of hydrogels over time under collagenase I. (G) Estimated degradation rates of hydrogels. Hydrogels become less degradable with addition of NVP or PEGDA. (H) FTIR spectra of hydrogels demonstrating incorporation of additional NVP and PEGDA, along with LAP photoinitiator. Individual data points indicate hydrogel replicates. Bars denote mean ± standard deviation. ** and *** indicate p<0.005 and p<0.001 respectively.

### 3.3 Hydrogel porosity

To test the effect of additional co-monomer incorporation on the resultant porosity of the hydrogel network, diffusion behavior of fluorescent TRITC-dextran was studied and quantified. Two different molecular weights of TRITC-dextran were tested, 4.4 kDa and 150 kDa (Figure 2B-E). For 4.4 kDa TRITC-dextran, release from the PF hydrogels was the fastest (significantly higher) when normalized to that in the other conditions (Figure 2B). However, due to its low hydrodynamic diameter (2.8 nm), the differences in diffusional release were not perceptible in the other hydrogel conditions. Hence, 150 kDa TRITC-dextran was used subsequently to obtain perceptible changes in normalized relative release from the hydrogel conditions (Figure 2C). For 150 kDa TRITC dextran, release was the quickest in PF conditions, followed by PF+5N and PF+10N, and being slowest in PF+1P and PF+2P conditions.

Based on the diffusion behavior, the diffusivity of TRITC-dextran was calculated for each hydrogel composition (Figure 2D). For 4.4 kDa TRITC-dextran, the diffusivity in PF hydrogels was calculated to be 2.6×10^-7^±2.9×10^-8^ cm^2^/s, while for the other hydrogel conditions, the average diffusivity value was 0.55×10^-7^±1.1×10^-8^ cm^2^/s. For the 150 kDa TRITC-dextran, the diffusivity in PF hydrogels was calculated to be 5.5×10^-8^±1.3×10^-9^ cm^2^/s. The average diffusivity in PF+5N and PF+10N hydrogels was 3.9×10^-8^±6.2×10^-9^ cm^2^/s and that in PF+1P and PF+2P hydrogels was 9.2×10^-9^±3.9×10^-9^ cm^2^/s (significantly lower). The theoretical mesh size of the hydrogel compositions was computed based on the diffusivity values of 150 kDa TRITC-dextran (Figure 2E). The mesh size of PF hydrogels was calculated to be 44.6±0.5 nm; the average mesh size of PF+5N and PF+10N hydrogels was 38.1±2.5 nm and that of PF+1P and PF+2P hydrogels was 24.2±3.0 nm (significantly lower) respectively.

These results demonstrate that incorporation of additional NVP in the base PF hydrogel slightly reduces the porosity of the hydrogel network due to additional crosslinks between the existing PEGDA chains and the NVP vinyl groups. However, incorporation of additional PEGDA in the base PF hydrogel significantly reduces the matrix porosity due to higher polymer content and presence of double networks within the hydrogel matrix.

### 3.4 Hydrogel degradability

To determine the relative enzymatic degradability of the hydrogel compositions, collagenase I at high activity (100U/mL) was used to monitor the degradation behavior over time (Figure 2F). PF hydrogels underwent complete degradation in 90 minutes, while PF+5N hydrogels underwent ∼57% degradation over 3 hours. The other hydrogel compositions displayed minimal degradation over time, as compared to the PEGDA control hydrogel. The degradation rate for these hydrogel compositions were calculated based on these time-dependent profiles (Figure 2G). PF hydrogels had a high degradability of 0.02±0.005 min^-1^, while PF+5N hydrogels had a degradability of 0.005±0.003 min^-1^ (significantly lower). The degradability rates of other hydrogels were negligible.

These results demonstrate the effect of incorporating a non-degradable co-monomer (NVP or PEGDA) within the base PF hydrogel composition. NVP, being low molecular weight (∼100 Da), has a gradual concentration-dependent decrease in degradability of the hydrogel network. Due to the presence of vinyl groups, the existing available acrylate groups of PF hydrogels form tighter low-degradable crosslinks, thereby resisting collagenase I action. However, addition of high molecular weight PEGDA (∼10 kDa) results in non-degradable double networks (PF-PEGDA and PEGDA-PEGDA), which restrict the accessibility of fibrinogen to the collagenase I. Although the degradation of the hydrogel network is partly mediated by the accessibility of the fibrinolytic cleavage sites and partly by the diffusion of collagenase I through the network, at high enzymatic concentrations, the proteolytic activity and hydrogel disassembly is expected to be much higher than diffusion-mediated transport rate of collagenase I.

### 3.5 Hydrogel chemical functionality

To further verify the incorporation of the non-degradable co-monomers within the hydrogel network, hydrogel samples were subjected to Fourier Transform Infrared (FTIR) spectroscopy (Figure 2H). The base PEGDA hydrogel showed a strong, sharp characteristic peaks at 1062-1098 cm^-1^ (C-O-C aliphatic ester), 1250-1480 cm^-1^ (C-H scissor and bending vibrations), and at 2870 cm^-1^ (CH_2_ stretching vibrations). Similarly, fibrinogen displayed characteristic peaks around 1630 cm^-1^, 3248 cm^-1^ (primary amine stretching). The peaks at 667 cm^-1^, 1646 cm^-1^, and 1735 cm^-^1 represent the C=C stretching and C=O stretching of the incorporated NVP. Additional PEGDA incorporation in base PF hydrogels was evident with the sharpening of the characteristic peaks for PEGDA. Overall, FTIR analysis exhibited successful incorporation and chemical modifications of the base PF hydrogel with additional co-monomers NVP or PEGDA.

### 3.6 Hydrogel ultrastructure

To visualize the ultrastructural morphology of the hydrogel compositions, scanning electron microscopy (SEM) was utilized (Figure 3). PF, PF+5N, and PF+10N hydrogel compositions (having low PEGDA content) showed a flaky, coarse morphology characteristic of the fibrinogen protein. The hydrogels also displayed a highly porous nature; however, no visible differences in porosity were observed among them. In comparison, PF+1P and PF+2P hydrogels (having higher PEGDA content) displayed a more uniform, denser, and less porous morphology. In particular, the relative appearance of pores in PF+2P hydrogels was significantly lower compared to other hydrogel conditions. PEGDA hydrogels (no protein content) displayed a dense, porous crosslinked network structure. Based on the PEGDA morphology, it can be concluded that PF+1P and PF+2P hydrogels display a double network crosslinked structure with the accessibility of the flaky fibrinogen structure being significantly reduced due to increased PEGDA content. These observations visually demonstrate the effect of additional PEGDA incorporation in PF hydrogels.

**Figure 3:**
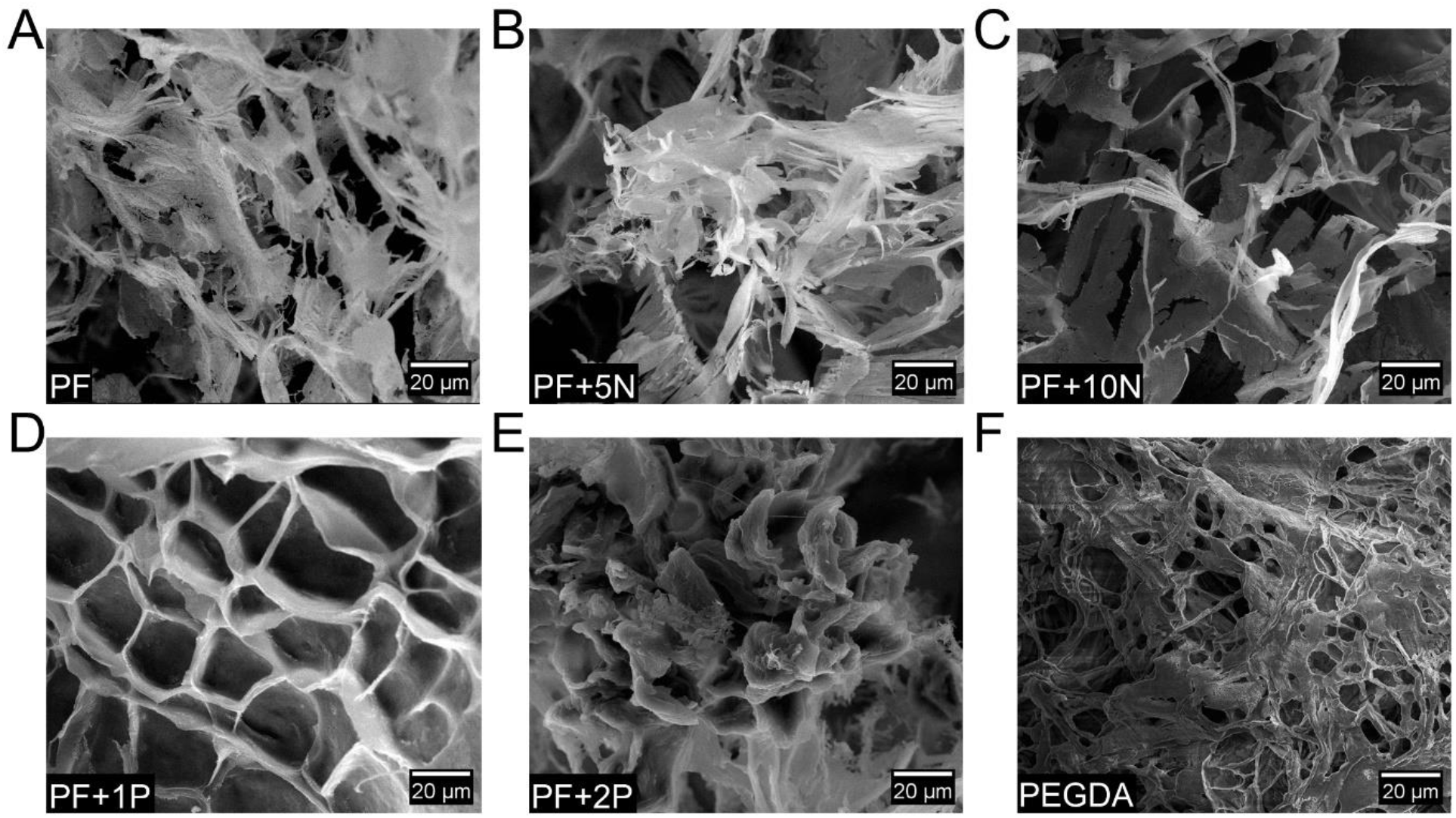
Hydrogel ultrastructure. (A-F) High magnification scanning electron micrograph images display the differences in hydrogel compositions. Fibrinogen appears as flaky morphology in (A-C). With additional PEGDA, hydrogels lose their flakiness and the porous, uniform morphology of the crosslinked PEGDA polymer becomes more prominent.

The porous morphology of the PF-based matrices visually observed through SEM indicate that the actual pore sizes are much larger than that computed through diffusion assay to obtain theoretical mesh sizes. The theoretical mesh sizes for the hydrogel composition vary between 20-50 nm, however, through rough visual inspection of SEM images, the pore sizes can reach up to 1 μm or even higher. These pore size dimensions would justify the ability of encapsulated cells to be able to migrate and spread through the matrix and their nuclei to be able to squeeze through the pores during the process.

### 3.7 Hydrogel mechanical properties

The effect of additional crosslinking of PF hydrogels via NVP or PEGDA on the mechanical properties of the hydrogel was assessed using rheometry (Figure 4). Under amplitude sweep (0.1 to 100% shear strain), the linear viscoelastic regions for the varying hydrogel compositions were determined. PF hydrogels showed a storage modulus of ∼30 Pa, loss modulus of ∼6 Pa and yield stress of 10 Pa (Figure 4Ai,ii,E). PF+5N and PF+10N hydrogels showed slightly increased moduli and yield stress (30 Pa and 75 Pa respectively), primarily due to more stable crosslinking of existing PEGDA polymeric chains. However, PF+1P and PF+2P hydrogels displayed significantly higher storage moduli (∼670 Pa and ∼1800 Pa respectively), loss moduli (∼30 Pa and ∼120 Pa respectively), and yield stress (330 and 980 Pa respectively), compared to other conditions, primarily due to additional crosslinking density of the excess PEGDA chains in the double network. Similarly, under frequency sweep (0.1 to 100 rad/s), PF hydrogels displayed the lowest storage and loss moduli, with additional NVP slightly enhancing the moduli (Figure 4Bi,ii). Additional PEGDA significantly increased the storage and loss moduli of the hydrogels, yielding similar results as the amplitude sweep. The estimated complex moduli, complex viscosity, loss tangent, and calculated compressive moduli (from the complex moduli) also showed similar trends indicating increased crosslinking and stiffness of the hydrogels with increasing PEGDA (Figure 4C-G). Loss tangent values of PF hydrogels were ∼0.1-0.2 and that of PEGDA hydrogels were ∼0.01-0.05 in the angular frequency range of 0.1-10 rad/s, with other hydrogel compositions having intermediate values (Figure 4E). The gel-sol transition of PF hydrogels was at 10 rad/s, while that of other hydrogels varied from 20-60 rad/s. Yield stress values of PF hydrogels was 10 Pa, which progressively increased with additional NVP or PEGDA to 980 Pa (Figure 4F). These values demonstrate differences in the viscoelastic properties and the gel-sol transition behavior of the hydrogels with additional NVP or PEGDA. The compressive moduli of the PF, PF+5N, PF+10N, PF+1P, PF+2P and PEGDA hydrogels was estimated to be 101±14 Pa, 566±64 Pa, 770±98 Pa, 1360±68 Pa, 4475±437 Pa and 1703±149 Pa respectively (Figure 4G). Overall, these results demonstrate the comparative role of additional NVP and PEGDA in contributing to the stiffness of PF hydrogels.

**Figure 4:**
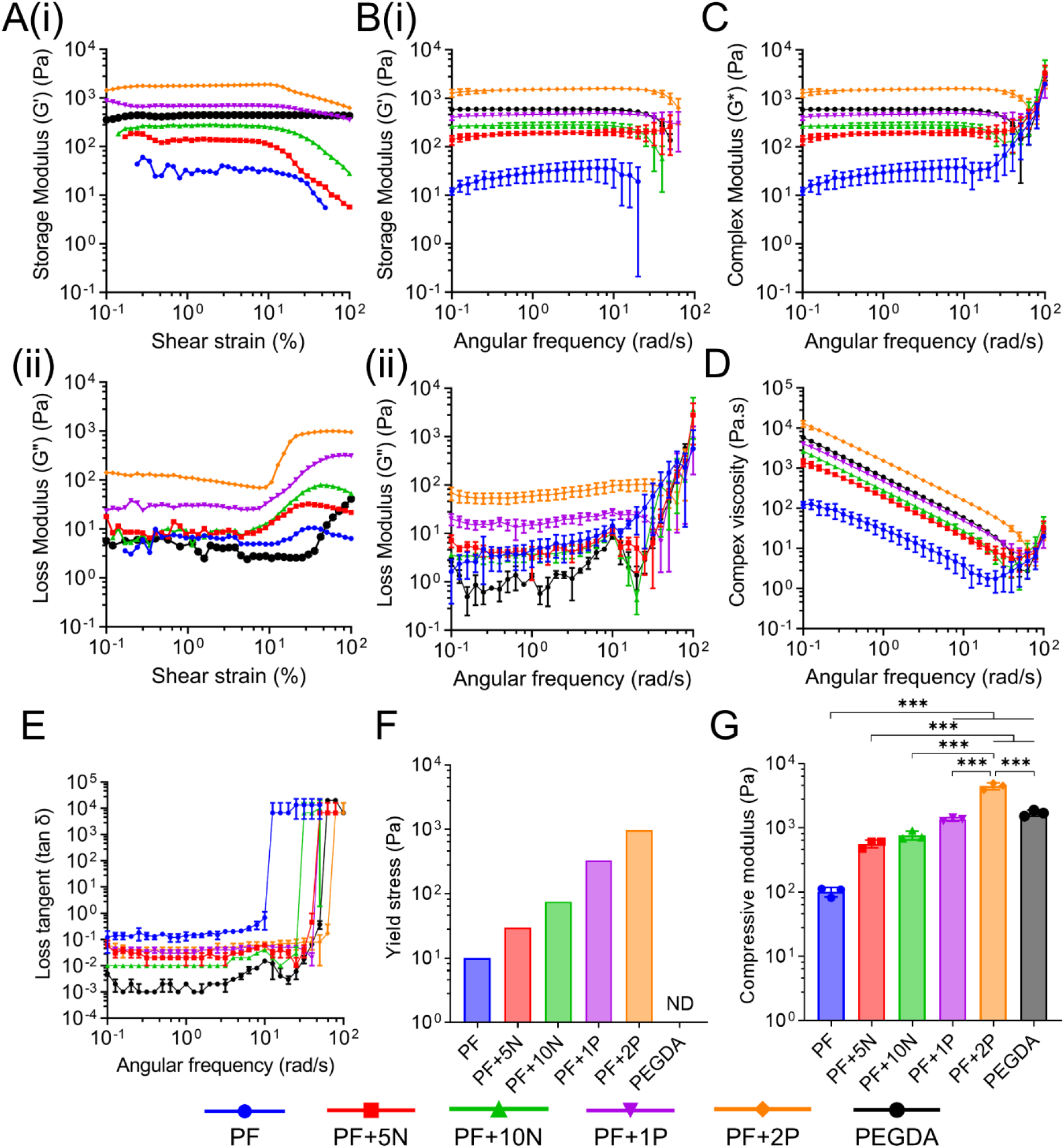
Rheological properties of hydrogels. (A) (i-ii) Storage and loss modulus of hydrogel compositions measured by amplitude sweep to determine linear range and yield point. (B) (i-ii) Storage and loss modulus of hydrogel compositions measured by frequency sweep. (C) Complex modulus, (D) complex viscosity, and (E) loss tangent estimated from frequency sweep. (F) Yield stress of hydrogel compositions measured from amplitude sweep. Hydrogel stiffness increases with additional NVP and further with additional PEGDA. Bars denote mean ± standard deviation. (A),(E) n=1 replicate. (B),(C),(D),(F),(G) n=3 replicates. *** indicates p<0.001.

The calculated complex moduli were further compared with estimated values from established theories of equilibrium swelling and rubber elasticity of polymer networks. The swelling behavior of hydrogels is based on osmotic pressure of ideal solution of polymer-solvent system. Similarly, the rubber elasticity theory is based on the assumption of Gaussian or ideal chain response for polymer segments. However, specifically designed and tunable hydrogels in this study have several factors which do not correspond to these ideal theories of rubber swelling and elasticity. Such factors include network topology, chain entanglements, semi-flexible nature of chains and electrostatic interactions. Due to end-crosslinking of PEGDA, multiple crosslinking points on the fibrinogen backbone, and copolymerization with NVP, there are distinct impacts on the nature of final network. Given the complex nature of network topology and chains between crosslink, the model proposed by Richbourg et. al. was adapted with some assumptions [25]. Details of the model parameters and equations are provided in Supplementary Methods. The model estimated values had good coherence with the experimentally obtained values from rheology (Supplementary Figure 1). For PF and PEGDA hydrogel compositions, the model values were ∼191% overestimated and ∼55% underestimated respectively. However, for the other hydrogel compositions, the model estimates were ∼12-22% of the experimental values, thereby demonstrating good reliability of the adapted model for application towards complex protein-network based polymeric hydrogels.

### 3.8 Fibroblast cellular and nuclear morphometry

To assess the effect of varying hydrogel composition and properties on 3D cellular behavior, NIH3T3 fibroblasts were cultured in 3D, and their morphology was assessed through fluorescence labeling and microscopy image analysis (Figure 5A-F). Cells in PF and PF+5N hydrogels displayed spread morphologies with elongated protrusions; this was significantly reduced in PF+10N, PF+1P, and PF+2P hydrogels, which featured more rounded cells. However, some cells displayed short protrusions in these hydrogels. Interestingly, cells in PEGDA hydrogels did not display any F-actin staining, possibly due to the absence of any adhesive ligands in the hydrogel matrix, and thereby no mechanoresponsive mechanism leading to polymerization of actin filaments. Cells were also uniformly distributed through the 3D imaged volume with their protrusions spreading primarily in a lateral direction, along with some axial spread (Supplementary Figure 2).

**Figure 5:**
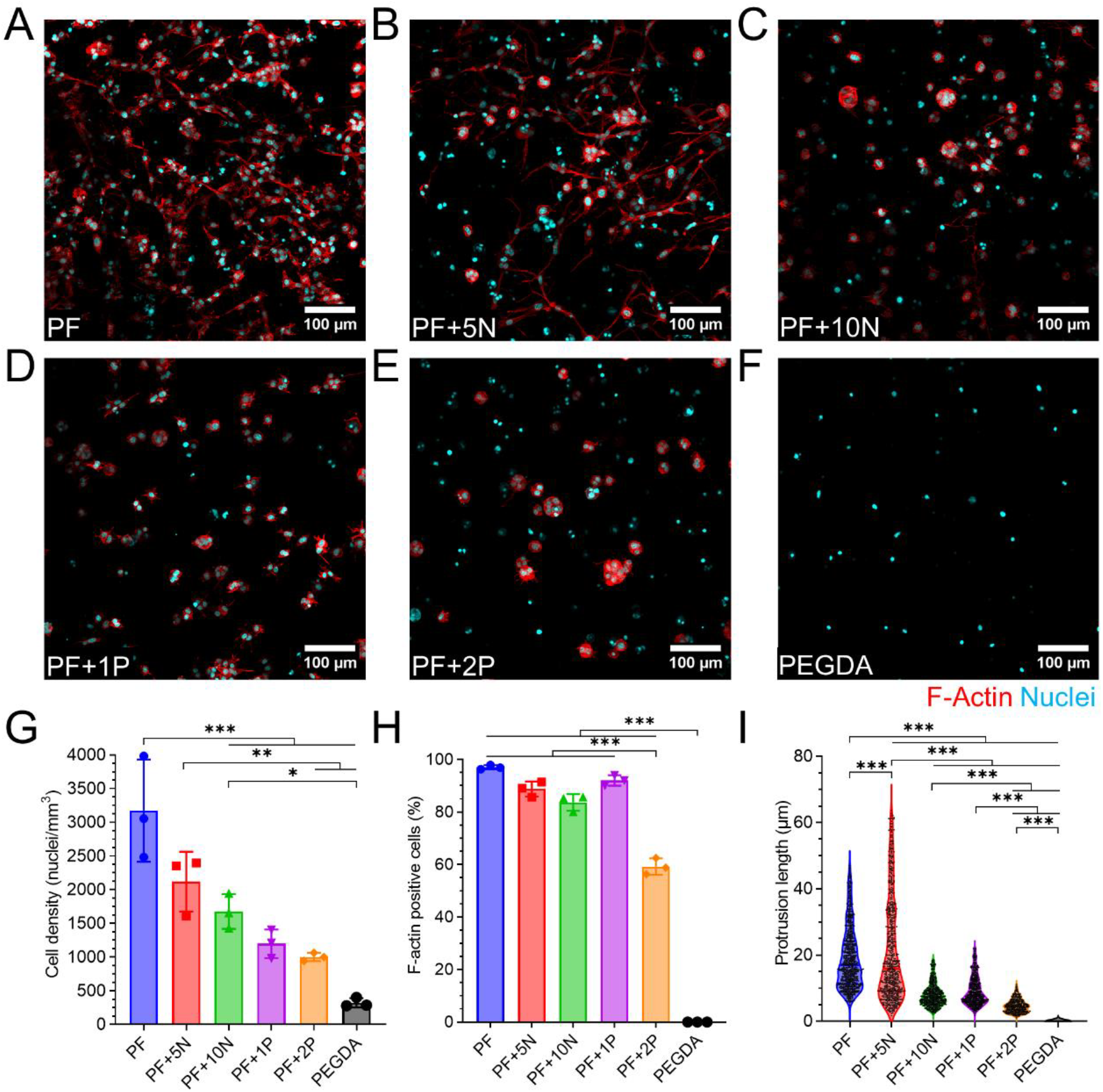
Fibroblast morphology and quantification. (A-F) Maximum intensity projections of 3D confocal z-stacks of NIH3T3 fibroblasts encapsulated in various hydrogel compositions on day 10 with labelled F-actin (Phalloidin, red) and nuclei (Hoechst 33342, blue). Cells in PEGDA hydrogels do not express F-actin. Scale bar=100 μm. (G) Cell density, (H) Percentage of F-actin positive cells, and (I) Protrusion length of cells measured from confocal z-stacks for various hydrogel compositions. (G-H) n=3 replicates. (I) Black points indicate individual protrusions (minimum=100) measured from at least 50 cells in each category. Bars denote mean ± standard deviation. *, **, and *** indicate p<0.05, p<0.005, and p<0.001 respectively.

Quantification of the cell density (number of nuclei per unit volume) revealed that PF hydrogels had the highest cell density (3172±758 cells/mm^3^), which progressively decreased with additional NVP and additional PEGDA in the PF matrix (Figure 5G). PF+2P hydrogels had a cell density of 1001±61 cells/mm^3^. Although the initial cell encapsulation density on day 0 was theoretically 5×10^6^ cells/mL (5000 cells/mm^3^), variations in cell density on day 10 could be attributed to cell spreading, proliferation, migration and possibly, varying degrees of equilibrium swelling of hydrogels.

The percentage of F-actin positive cells were quantified in each hydrogel composition with PF, PF+5N, PF+10N, and PF+1P hydrogels displaying comparable percentages (>85% on average), indicating good cellular adhesion with the hydrogel matrix (Figure 5H). However, cells in PF+2P hydrogels displayed only ∼59% F-actin positive cells, indicating restricted adhesion of cells to the matrix possibly due to hindered accessibility of the fibrinogen adhesive moieties by the PEGDA chains.

The protrusion length and protrusion frequency (number of protrusions per cell) were quantified for cells in each hydrogel composition (Figure 5I, Supplementary figure 3A). Cells in PF and PF+5N hydrogels displayed comparably high protrusion lengths (19±9 μm and 21±15 μm respectively). In comparison, cells in PF+10N, PF+1P and PF+2P displayed significantly reduced protrusion lengths (8±3 μm, 9±4 μm, and 5±2 μm respectively). In PF, PF+5N, PF+10N, and PF+1P hydrogels, cells displayed on average 2-3 protrusions per cell; however, in PF+2P hydrogels, cells displayed on average <1 protrusion per cell, thereby indicating the restrictive nature of the PF+2P hydrogels. Overall, these measurements provide an estimate of the restrictive effect of additional NVP and PEGDA on 3D fibroblast cell density, cell adhesion to the matrix, and cell spreading.

The cellular and nuclear morphologies of the fibroblasts (area, diameter, circularity, and aspect ratio) were further quantified as a function of each hydrogel composition (Figure 6). PF and PF+5N hydrogels had higher average projected cell area (405±150 μm^2^) compared to PF+10N, PF+1P, and PF+2P hydrogels (289±108 μm^2^) (Figure 6A). In PEGDA hydrogels, due to absence of F-actin staining, the cellular morphological parameters could not be determined (Figure 6A-D). PF and PF+5N hydrogels also displayed higher mean geometric cell diameter (22±4 μm vs. 19±3 μm), lower circularity (0.39±0.19 vs. 0.65±0.53) and higher aspect ratio (3.1±1.5 vs. 1.5±0.4) compared to that in PF+10N, PF+1P and PF+2P hydrogels. These results collectively indicate that addition of low concentration of NVP (5 mM) does not restrict cell spreading and elongation compared to PF hydrogels. In fact, owing to the increased stability of the crosslinked hydrogels, cells in PF+5N hydrogels were observed to be more spread out than in PF hydrogel condition. However, with higher NVP concentration (10 mM) or additional PEGDA in the matrix, the cells are restricted from spreading and remain circular in shape, unable to form elongated morphologies.

**Figure 6:**
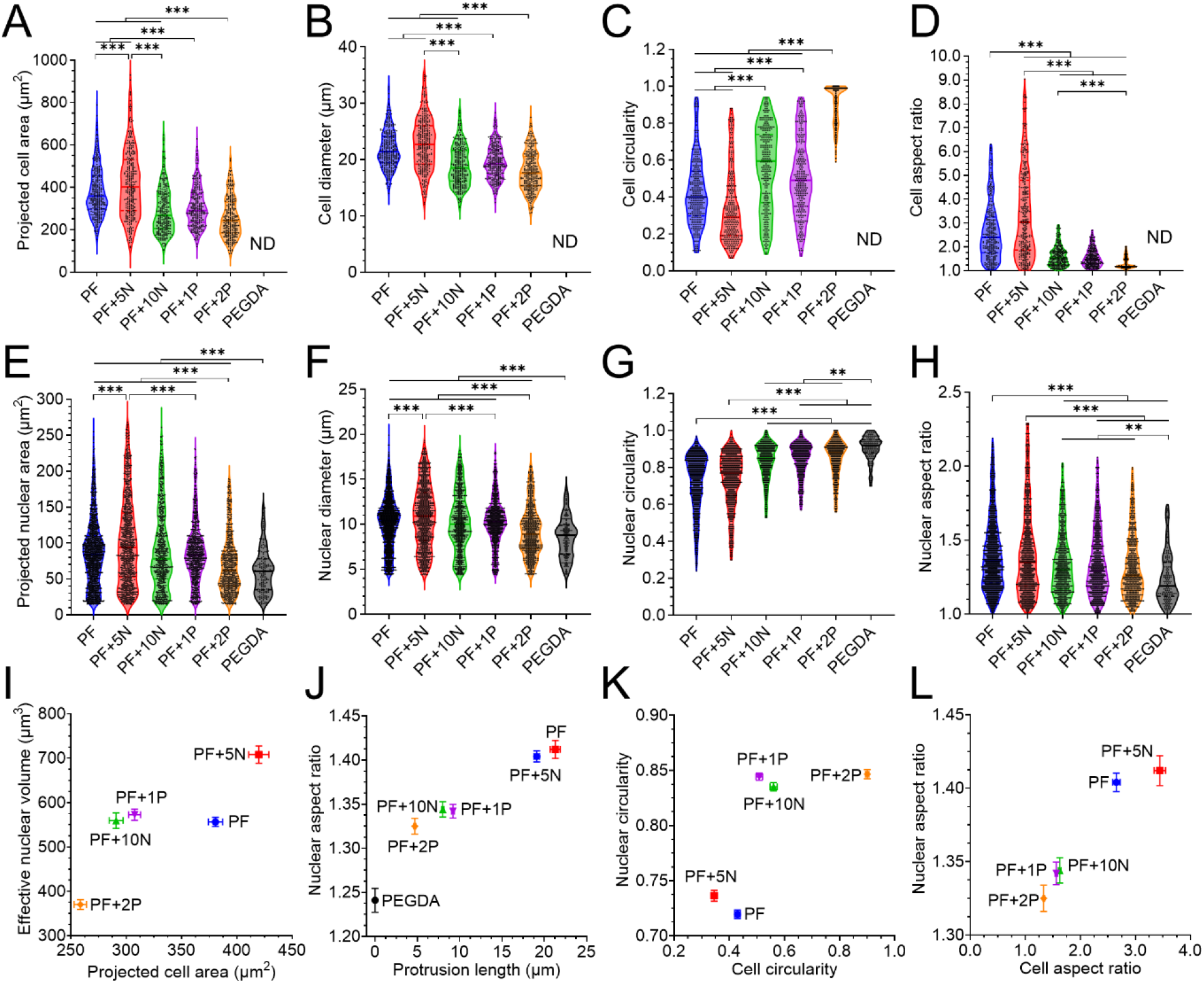
Cellular and nuclear morphometry of fibroblasts. (A) Projected cell area, (B) mean geometric cell diameter, (C) cell circularity, (D) cell aspect ratio of NIH3T3 fibroblasts measured on day 10 of culture. (E) Projected nuclear area, (F) mean geometric nuclear diameter, (G) nuclear circularity, and (H) nuclear aspect ratio of NIH3T3 nuclei. Each black dot represents an individual cell (A-D) or nuclei (E-H). n=minimum 200 cell or nuclei for each category. ** and *** indicate p<0.005, and p<0.001 respectively. Cellular and nuclear morphological parameters significantly change with additional NVP or PEGDA. (I) Correlation plot of effective nuclear volume vs. projected cell area, (J) nuclear aspect ratio vs. protrusion length, (K) nuclear circularity vs. cell circularity, and (L) nuclear aspect ratio vs. cell aspect ratio for each category. Values denote mean ± standard error. Hydrogel compositions with less confined cells appear as a distinct cluster than those with more confined cells.

Similar to cellular morphology, nuclear morphology also varied with hydrogel composition, albeit to varying degrees (Figure 6E-H). Projected nuclear area in PF hydrogels did not decrease with additional NVP or PF+1P (92±51 μm^2^) but decreased significantly only in the PF+2P and PEGDA hydrogels (66±36 μm^2^). Similarly, nuclear diameter was significantly reduced in PF+2P and PEGDA hydrogels compared to other conditions (8.9±2.5 μm vs.10.4±3.1 μm). Nuclear circularity and aspect ratio (indicative of the nuclear deformation during cell spreading) was significantly different in PF and PF+5N hydrogels compared to other conditions (0.73±0.15 vs. 0.86±0.09 and 1.4±0.3 vs. 1.3±0.2 respectively). The effective nuclear volume (indicative of nuclear confinement) was also significantly lower in PF+2P and PEGDA hydrogels compared to other conditions (345±230 μm^3^ vs. 600±445 μm^3^) (Supplementary Figure 3B). These results indicate that the effect of matrix composition and properties are translated to not only cellular morphological features, but to nuclear morphological features as well.

The translational effect of cellular morphology on nuclear morphology was further investigated by correlating specific measured parameters (Figure 6I-L). For each hydrogel condition, the effective nuclear area was correlated against projected cell area (Figure 6I). PF+5N hydrogel and PF+2P hydrogels were the most distinct and distant groups in the correlation plot, with other hydrogel compositions lying in between. This indicates that cells with higher area (highly spread cells) in PF+5N also have larger nuclei, while cells in PF+2P hydrogels which are more confined have smaller nuclei in comparison. Similarly, the nuclear aspect ratio and nuclear circularity (indicative of nuclear deformation during cell migration) was correlated with protrusion length, cell circularity and cell aspect ratio (indicative of cell spreading and 3D migration). PF and PF+5N hydrogels formed a distinct cluster compared to other hydrogel compositions, further quantitatively validating the fluorescence z-stack observations that these two hydrogel conditions allowed higher cell spreading and elongation, better cell-cell connectivity and attachment, compared to other hydrogel conditions, where cells were mostly singular, rounded and confined in their local niche.

### 3.9 Effect of matrix compliance on cellular and nuclear confinement

The matrix properties measured for each of the hydrogel compositions was further correlated to the cellular and nuclear morphological parameters, to develop a holistic understanding of the cellular response to hydrogel matrix (Figure 7). The four primary matrix properties, adhesivity (measured by fibrinogen content), degradability (measured by collagenase I enzymatic activity), porosity (measured by TRITC-dextran diffusivity and theoretical mesh size), and stiffness (measured by compressive modulus from rheology), were normalized and scored between 0.0 and 1.0 (indicating lowest and highest values). The cellular and nuclear morphological features measured earlier were also normalized between 0.0 and 1.0 and correlated with the matrix properties (Figure 7A). The correlation heatmap provides a comprehensive summary of the various observations and provides perspective on the individual role of each parameter towards specific features of the cell and nucleus.

**Figure 7:**
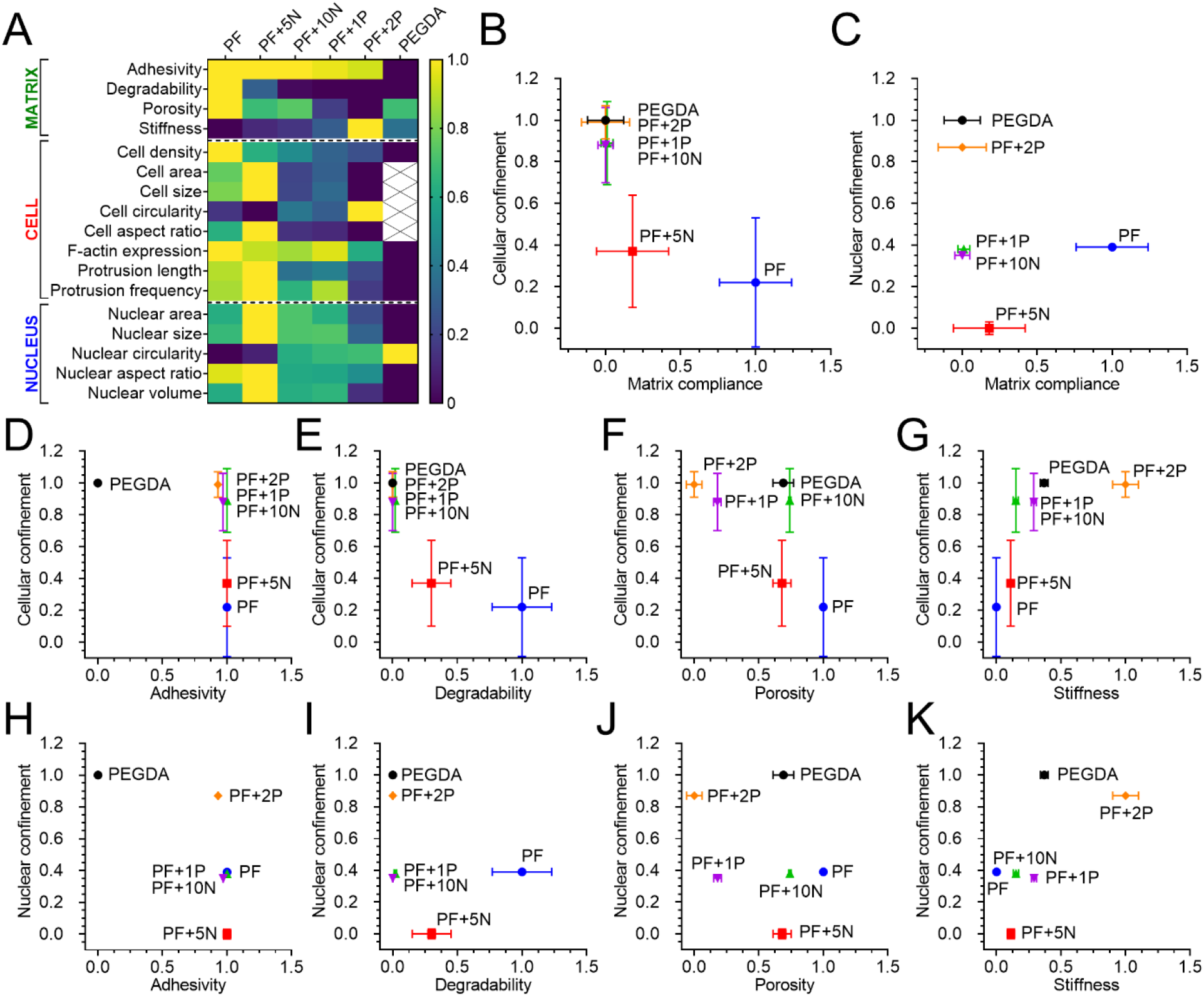
Role of matrix compliance on cellular and nuclear confinement. (A) Correlation heatmap of various matrix properties, and cellular and nuclear morphological features across various hydrogel compositions. Values are normalized and color-coded between 0.0 and 1.0. (B-C) Correlation plot of cellular and nuclear confinement as a function of matrix compliance. Correlation plot of (D-G) cellular confinement and (H-K) nuclear confinement vs. matrix adhesivity, degradability, porosity, and stiffness. Values denote mean ± standard error (of normalized means). Distinct clusters emerge among the various hydrogel compositions based on individual matrix properties.

Based on the correlation heatmap, the four defining matrix properties (adhesivity, degradability, porosity, and stiffness) were collectively termed as ‘matrix compliance’. Similarly, the cellular features (cell density, protrusion length, and protrusion frequency) were collectively termed ‘cellular confinement’, and the effective nuclear volume as termed ‘nuclear confinement’. PF hydrogels having the highest fibrinogen content, highest degradability to collagenase I, highest diffusivity and mesh size, and lowest stiffness was considered to have the highest matrix compliance of 1.0, while PEGDA hydrogels with zero fibrinogen content, no response to collagenase I enzymatic activity, and medium diffusivity and mesh size, and medium stiffness was considered to have the lowest matrix compliance of 0.0. High cell density and high protrusivity indicated low cellular confinement (0.0), while low cell density and low protrusivity indicated high cell confinement (1.0). Similarly, high nuclear volume indicated low nuclear confinement (0.0) and low nuclear volume indicated high nuclear confinement (1.0).

Using this correlative classification technique, it was observed that PF+10N, PF+1P, PF+2P, and PEGDA hydrogels formed a distinct cluster with respect to cellular confinement (high values close to 1.0), which corroborated the initial observations in confocal z-stack images (Figure 7B). Additionally, PF+2P and PEGDA hydrogels formed a distinct cluster with respect to nuclear confinement (high values close to 1.0). Correlation of cellular confinement and nuclear confinement with matrix adhesivity, degradability, porosity, and stiffness also provided a rough estimate of the contribution of individual matrix features ((Figure 7D-K). Further deconvolution and correlation of cellular confinement (cell density, protrusion length, and protrusion frequency) with individual matrix features further corroborated these analyses (Supplementary Figure 4). In all comparisons, PF+2P hydrogel condition was observed to have high degree of cellular and nuclear confinement, consistent with its highly restrictive features, and closely matching that of the control PEGDA hydrogel condition. PF and PF+5N hydrogels were most compliant and low confining towards cells. Overall, this approach provides a streamlined approach to correlate the role of specific matrix features with cellular and nuclear morphological features, thereby providing insights into biological response in functional and tunable engineered matrices.

## 4. DISCUSSION

Hydrogels are versatile materials for application towards tissue engineering, regenerative medicine, and general investigations of cell-biomaterial interactions. To that end, the biochemical and biophysical properties of hydrogels that affect 3D cellular behavior is of particular interest to researchers. Among various properties, the role of matrix stiffness on 3D cell behavior and function has received wide attention, primarily owing to differences between physiological and pathological tissue microenvironments in the context of cancer, aging, fibrosis and others [26,27].

In 2D culture studies, where cells are seeded on top of flat substrates, the role of matrix stiffness on cell behavior and function has been studied extensively [28,29]. In such systems, it is fairly easy to decouple the effect of matrix adhesivity and porosity from matrix stiffness on cell spreading and/or differentiation as substrates can be engineered in a controlled manner. Hence the relative contribution of each matrix feature towards specific cell behavior can be estimated and major biophysical/biochemical drivers of pathological progression in the ECM can be identified. In 3D culture studies, the matrix factors influencing cell behavior (adhesivity, degradability, porosity, stiffness, fiber alignment, anisotropy) become compounded and coupled to each other, making it difficult to delineate their role or relative contribution to cell behavior.

One standard approach to mitigate this challenge is to incorporate specific functional ligands to hydrogel matrices independent of each other [30,31]. For example, incorporation of RGDS peptide sequence into hydrogel matrices can control matrix adhesivity independent of other features (e.g., stiffness, degradability, porosity etc.) [32]. Similarly, incorporation of proteolytic peptide sequence into hydrogel matrices can control degradability independent of matrix adhesivity, porosity, and stiffness [33]. However, this systematic approach is difficult to implement in complex protein– and polysaccharide-based hydrogels, as the proteolytic sites and adhesive ligands are often present in the same polymer backbone constituting the hydrogel. In addition, the network functionality of the same polymer backbone also dictates the degree of crosslinking which in turn influences the porosity and stiffness of the resulting hydrogel matrix. Hence, it becomes important to simultaneously correlate multiple defining matrix features with cellular (and nuclear) behavior to obtain a holistic understanding of the cell-material response.

In this study, PEGylated fibrinogen is used as the base polymer to investigate the behavior of fibroblasts through cellular and nuclear morphometry. The fibrinogen backbone contains both the adhesive and proteolytically cleavable sites that confer matrix adhesivity and degradability to the resulting hydrogels. When fibrinogen is covalently conjugated with PEGDA (using one of the acrylate end groups) and the resulting PF hydrogel is crosslinked, the dangling acrylate end groups of the PEGDA chains provide the template to form additional crosslinks that ultimately influence the crosslinking density (and hence porosity and stiffness) of the hydrogel network. Additionally, considering that fibrinogen has significantly higher molecular weight (340 kDa) than PEGDA (10 kDa), it is expected that fibrinogen by itself would also contribute significantly to the overall stiffness and porosity in the final PF hydrogel matrix. Overall, in the PF hydrogel network, the matrix adhesivity and degradability is attributed to fibrinogen content, while matrix porosity and stiffness is attributed to both fibrinogen and PEGDA constituents.

Modifications to the based PF hydrogel network has been achieved via two means: 1) addition of a non-degradable, ‘zero-length’ co-monomer NVP, and 2) addition of a non-degradable, long chain co-monomer PEGDA. In the first case, NVP by itself lacks the ability to form independent hydrogel networks, but forms stronger crosslinks and reinforces the existing dangling PEGDA side chains in the PF hydrogel network by forming an intermediate adduct poly(ethylene glycol acrylate-co-vinyl pyrrolidone) [34]. However, this mechanism of crosslinking is limited by the available number of free acrylate groups in the dangling PEGDA chains. In this study, 5 mM and 10 mM NVP were chosen (representing a molar ratio of NVP:free available acrylate of 7:1 and 14:1 respectively), as higher NVP concentrations is expected to get saturated and remain unutilized in a limited amount of PEGDA acrylates. The additional NVP does not contribute significantly to the total polymer content (due to its low molecular weight). However, it significantly reduces the flexibility of the crosslinked PEGDA chains, forming tighter and stabler junctions, leading to decreased swelling capacity, slight reduction in porosity (theoretical mesh size) and moderate increase in stiffness. However, there is a significant reduction in matrix degradability with additional NVP, thereby indicating the strengthening and stability of the crosslinked network. In the second case, addition of excess PEGDA leads to increase in total polymer content, total number of free acrylates available for additional crosslinking, thereby leading to a double interpenetrating network consisting of PF-PEGDA and PEGDA-PEGDA crosslinks. This network is characterized by significantly higher crosslinking density, leading to decreased porosity and increased stiffness compared to the base PF matrix. In addition, since PEGDA is non-degradable in nature, the additional crosslinks reduce the accessibility of the adhesive and proteolytic sites on the fibrinogen backbone, thereby reducing degradability and adhesivity of the PF+1P and PF+2P matrices significantly.

The effects of these matrix modifications are manifested through changes in the cellular and nuclear morphology of encapsulated fibroblasts. In PF hydrogels (highly compliant and well permissive matrix), fibroblasts display elongated and well spread morphologies with abundance of filopodial protrusions. Even in PF+5N hydrogels which have significantly reduced degradability compared to PF, fibroblasts were observed to be well-spread and connected to each other, thereby indicating that these cells do not depend solely on proteolytic degradation for 3D migration but can switch from mesenchymal to amoeboid mode of migration for spreading. In fact, the improved stability of the PF+5N matrix compared to the PF matrix may even lead to enhanced cell migration and spreading, as evidenced longer and denser protrusions of the cells. However, beyond a certain limit (10 mM NVP in this case), the overall compliance of the matrix is significantly reduced, and the cells become confined and restricted. Similar observations are also validated in PF+1P and PF+2P with low matrix compliance. Particularly, cells encapsulated in PF+2P showed variations in their morphology. Encapsulated cells which had access to the adhesive sites of fibrinogen displayed F-actin expression and small protrusions, however, significant number of cells which had restricted access to the fibrinogen adhesive sites displayed no F-actin expression and hence no protrusions.

The effects of the cellular deformation (spreading vs. confinement) were also transmitted to the nuclear level. Well-spread cells in PF and PF+5N hydrogels displayed elongated, larger nuclei indicating that the nuclei were also able to undergo deformation (owing to matrix being compliant). Particularly, in PF+5N hydrogels, nuclear confinement was observed to be the lowest, indicating that the nuclei of the encapsulated cells were able to squeeze through the matrix pores during cell migration and spreading. However, in other low-compliant (PF+10N, PF+1P, PF+2P) and non-compliant (PEGDA) matrices, the nuclei appeared more rounded and restricted.

In previous studies, various 3D hydrogel systems have been used to investigate the effect of matrix properties (stiffness, degradability) on fibroblast morphology and spreading [10,20,31,35,36]. The effect of matrix stiffness and proteolytic degradation on smooth muscle cell migration has been partly uncoupled [37]. Although most literature in the field attribute morphological changes in cell behavior primarily to matrix stiffness, the role of other matrix features, namely, adhesivity, porosity, and degradability also play influential roles in 3D cell migration and spreading [38,39]. Controlled investigations of nuclear deformation under varying levels of 2D and 3D confinement within interconnected microfluidic compartments have also revealed specific mechanisms of cancer cell migration, particularly in the context of mesenchymal-amoeboid mode of transition while navigating tight cellular barriers [40,41]. The nuclear deformability of the cell may be primarily governed by the matrix porosity; however, matrix stiffness and degradability may also play influential roles, particularly over longer time scales of 3D cell culture studies. In this study, it is challenging to completely decouple the contribution of individual matrix features to cell and nuclear phenotype, as the adhesive and proteolytic sites are encoded in the same fibrinogen backbone. Also, addition of NVP or PEGDA affects both degradability and stiffness to varying levels. In the future, independent peptide sequences having adhesive or proteolytic sites could be used to have independent control of these features. Nevertheless, the presented approach provides some insight towards partly decoupling the effects of individual matrix features on cellular and nuclear morphogenesis and 3D behavior.

## 5. CONCLUSIONS

This study provides insights into cellular and nuclear deformation and 3D response of fibroblasts to varying matrix conditions, collectively termed ‘matrix compliance’ and decoupled into four defining features: adhesivity, degradability, porosity, and stiffness. PF-based hydrogels were selectively modified by addition of two different non-degradable co-monomers of varying molecular weight (0.1 kDa NVP or 10 kDa PEGDA), thereby leading to two different modes of crosslinking and resulting in hydrogels with varying degrees of adhesivity, degradability, porosity, and stiffness. Cellular and nuclear morphometry revealed that PF and PF+5N (PF modified with 5 mM NVP) hydrogels had high matrix compliance and permitted 3D cell spreading, high protrusivity, higher cell density, and associated nuclear elongation. However, PF+2P (PF modified with 2% w/v PEGDA) had low matrix compliance and restricted cell spreading, formation of protrusions, low cell density with confined rounded nuclei. These observations provide insights into 3D cell-material response, which can be implemented in the future for intelligent design of biomaterial or hydrogel matrices, wide range of tissue engineering applications, modeling of pathological phenomena, or investigation of structure-function relationships in 3D morphogenesis.

## 6. SUPPLEMENTARY MATERIAL

Supplementary figures and Supplementary Methods are available in the Appendix.

## 7. ACKNOWLEDGEMENTS

This study was supported by funding from DBT Ramalingaswami Fellowship, Department of Biotechnology, India (SP) (BT/RLF/Re-entry/16/2019), SERB Startup Research Grant (SP) (SRG/2020/000343), and Indian Institute of Technology Madras Seed Grant (SP). The authors acknowledge support from Sophisticated Analytical Instrument Facility (SAIF), IITM for SEM imaging. The confocal microscopy facility is supported by the DST-FIST Advanced Bioimaging Facility (SR/FST/LS-II/2020/552(C)) and Industrial Consultancy and Sponsored Research (ICSR)-Common Instruments Facility (CIF), IIT Madras. IPP is supported by UGC Fellowship, Ministry of Human Resource Development (MHRD), India and Women Leading IITM Fellowship, IIT Madras.

## 9. CRediT AUTHORSHIP CONTRIBUTION STATEMENT

IPP: Hydrogel characterization, cell studies, and manuscript writing. SS (Saujanya): Hydrogel characterization and theoretical modeling. SS (Shiuly): Hydrogel characterization. AD: Supervision on hydrogel characterization and theoretical modeling. SP: Conceptualization, design and supervision, image analysis, data analysis, and manuscript writing. All authors provided critical feedback and helped shape the research, analysis, and the manuscript.

## APPENDIX

### Supplementary Figures

**Supplementary Figure 1:**
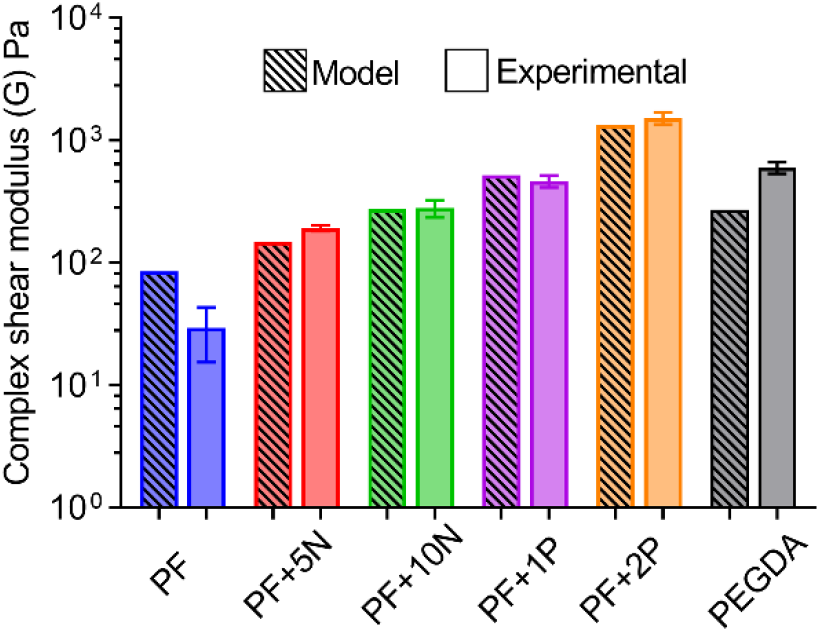
Comparison of complex shear modulus as obtained from the model vs. experimental values obtained from rheology (data represented as mean ± standard deviation of three hydrogel replicates with their complex modulus measured by frequency sweep at 1 rad/s)

**Supplementary Figure 2:**
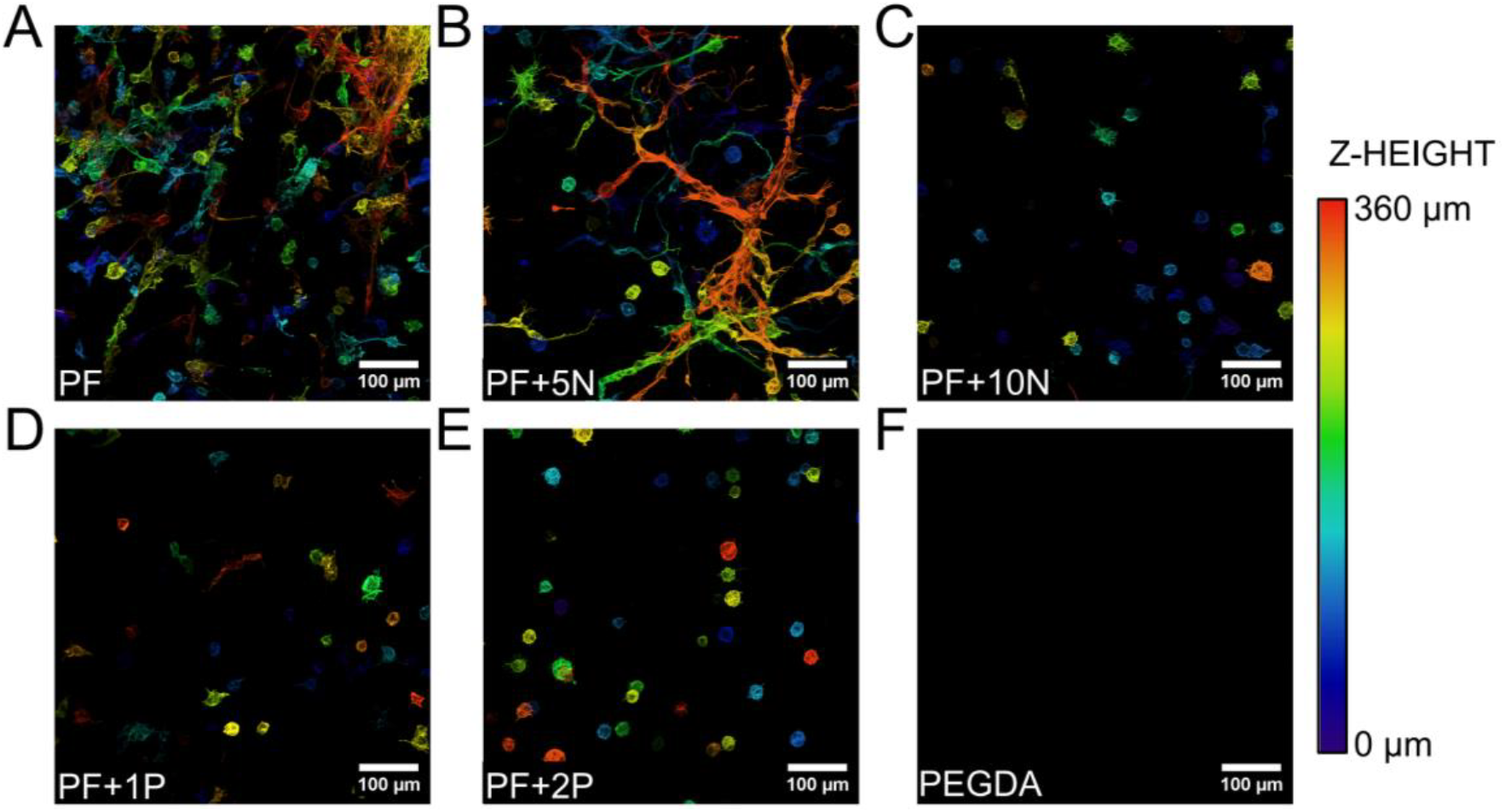
Cellular morphology of fibroblasts in tunable hydrogels. NIH3T3 fibroblasts encapsulated in (A) PF, (B) PF+5N, (C) PF+10N, (D) PF+1P, (E) PF+2P, and (F) PEGDA hydrogels, fixed and stained for F-actin (phalloidin) on Day 10 and z-depth color-coded to represent the distribution and spread within a 3D volume of depth 360 μm.

**Supplementary Figure 3:**
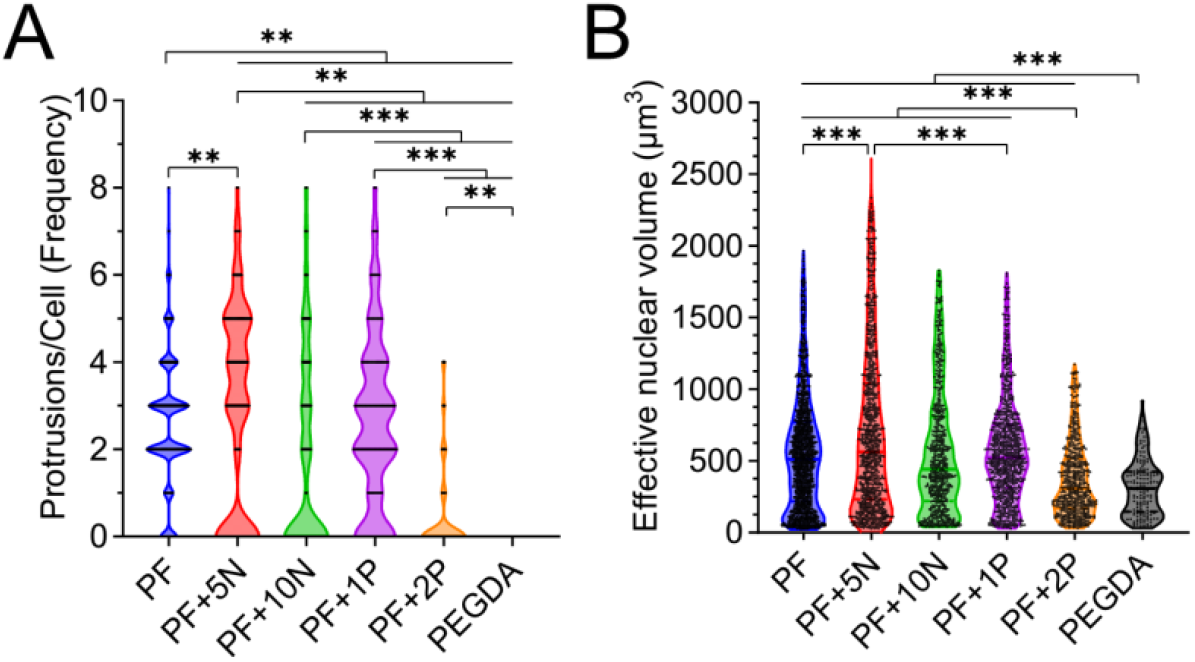
Cellular and nuclear morphology of fibroblasts. NIH3T3 fibroblasts stained for F-actin (phalloidin) and nuclei (Hoechst 33342) and imaged and analyzed for (A) average number of cellular protrusions per cell, and (B) calculated effective nuclear volume, within different hydrogel conditions. Each black dot represents an individual cell (A) or nuclei (B). Median and upper and lower quartiles are indicated in the violin plots. n>100 cells or nuclei per category. **, and *** indicate p<0.005, and p<0.001 respectively.

**Supplementary Figure 4:**
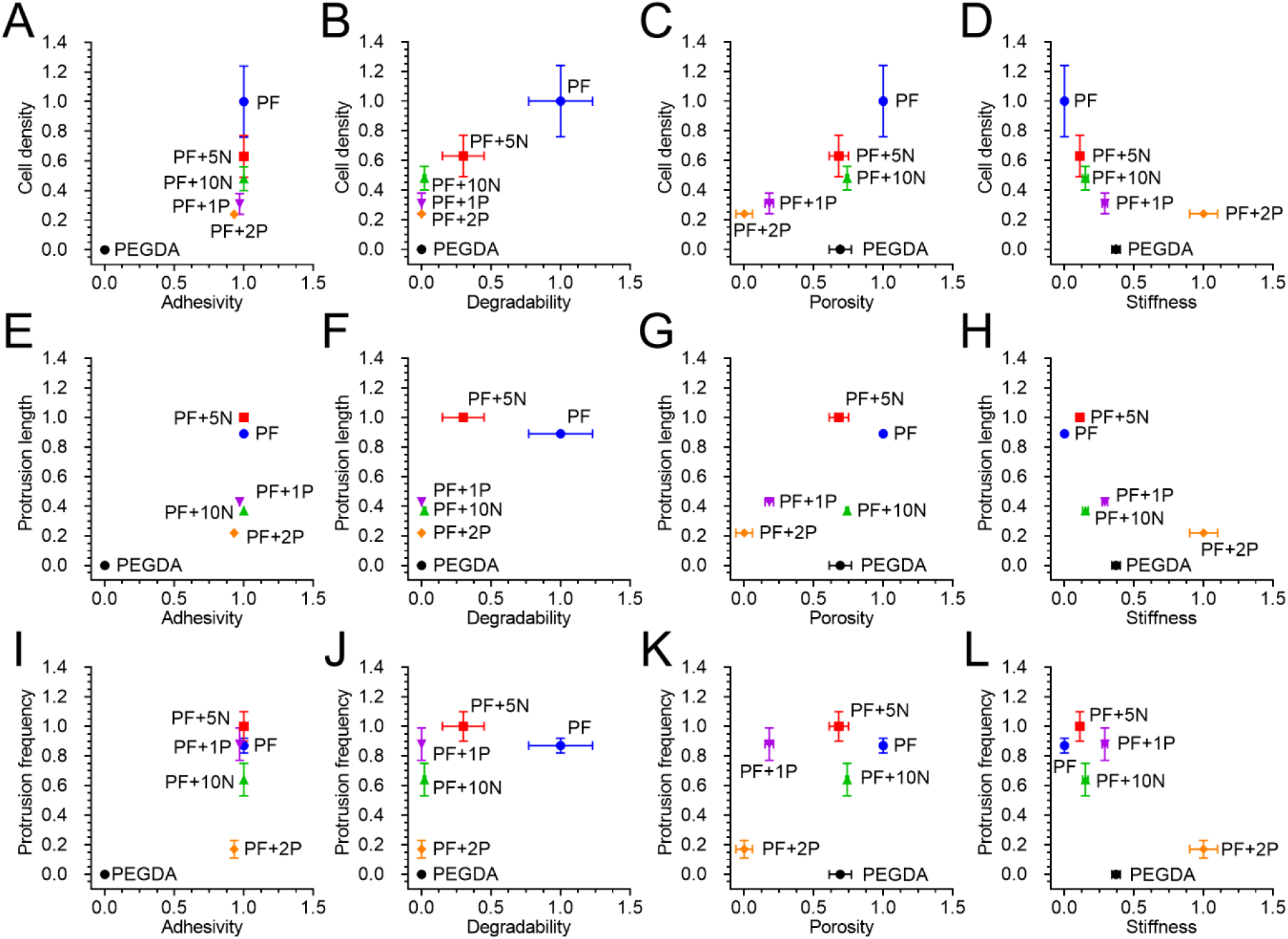
Cellular morphology correlated with matrix properties. (A-D) Normalized cell density, (E-H) normalized protrusion length, and (I-L) normalized protrusions per cell in 6 hydrogel conditions correlated with matrix adhesivity, degradability, porosity, and stiffness. Data represented as mean ± standard error (of the normalized values).

### Supplementary Methods

All chemicals were obtained from Sigma-Aldrich unless otherwise mentioned.

#### 1. PEGDA synthesis

Poly(ethylene glycol) (PEG) (molecular weight: 10 kDa) was reacted with acryloyl chloride at a molar ratio of 1:4 in anhydrous dichloromethane with triethylamine (TEA) (molar ratio of PEG:TEA = 1:2) under argon overnight at 25°C. Post-reaction, 1.5M K_2_CO_3_ was added to the reaction mixture to purify the resultant PEGDA via phase separation. The organic phase, containing PEGDA, was dried over anhydrous MgSO_4_ to remove residual aqueous content, followed by filtration. PEGDA was then precipitated in 4 L of cold diethyl ether, collected by filtration, and vacuum-dried overnight at 25 °C. The final product was stored at –20 °C as a dry powder.

#### 2. PEG-fibrinogen synthesis

Bovine fibrinogen was dissolved in 8M urea in 10 mM PBS (pH 7.4) at a concentration of 7 mg/mL. To facilitate reduction, tris(2-carboxyethyl) phosphine hydrochloride (TCEP-HCl) was added at a molar ratio of 1:5:1 (TCEP to fibrinogen cysteines), and the final pH was adjusted to 8.0. Separately, PEGDA (synthesized earlier) was dissolved in 8M urea-PBS at a concentration of 280 mg/mL and gradually added to the fibrinogen solution. The reaction was allowed to proceed under dark for 3 hours at 25°C. To purify the PEGylated fibrinogen from the unreacted PEGDA, the reaction mixture was diluted with an equal volume of urea-PBS buffer and precipitated by adding acetone at a 4:1 volumetric ratio (acetone to product solution). The precipitate was then separated by centrifugation, carefully collected, and weighed. For complete dissolution, the precipitate was resuspended in urea-PBS buffer at a ratio of 2 mL of buffer per gram of precipitate, followed by thorough homogenization. The solution underwent dialysis against 4 L of sterile PBS for 24 hours at 4°C, with three buffer changes. The purified PEG-Fb product was aliquoted into sterile centrifuge tubes and stored at –80°C for long-term preservation.

#### 3. LAP synthesis and characterization

Dimethyl phenylphosphonite was reacted with 2,4,6-trimethylbenzoyl chloride in an equimolar ratio under inert argon and allowed to react overnight for 18 hours under continuous stirring. Four-fold excess lithium bromide was dissolved in 2-butanone, added to the reaction mixture and then heated to 50°C. The solid precipitate formed after 10 minutes was cooled to ambient temperature, allowed to rest for 4 hours, and filtered. The filtrate was washed and filtered with 2-butanone three times to remove unreacted lithium bromide. The resulting solvent was left to evaporate for a few hours and further removed by freeze-drying. The obtained product lithium phenyl-2,4,6-trimethylbenzoyl phosphinate (LAP) was stored in argon-filled vials at 4°C under dark. The completion of the reaction of the LAP product was verified by ^1^H NMR spectroscopy.

#### 4. Diffusion and porosity estimation

Briefly, photocrosslinked hydrogels (4 mm diameter and 1 mm thickness) were swollen to equilibrium and incubated overnight in TRITC-dextran (molecular weight 4.4 kDa or 150 kDa, Thermo Fisher Scientific) in PBS at 0.5 mg/mL to ensure equilibrium loading. Following incubation, the TRITC-Dextran solution was completely removed from each well and replaced with 200 μL PBS, marking the start of the release study. At 15-minute intervals, over a 3-hour period, 50 μL of solution was collected from each well and replaced with an equal volume of fresh PBS buffer. The fluorescence intensity of the released TRITC-Dextran was measured using a plate reader (Biotek Synergy H1, excitation: 540/25 nm, emission: 590/35 nm). A standard curve was generated using known TRITC-Dextran concentrations in PBS, while fresh PBS served as a blank control. A minimum of 5 replicates per condition were measured. The cumulative mass release of TRITC-Dextran was analyzed from fluorescence intensities, and the diffusion coefficient (D) was determined using the following Equation (1).

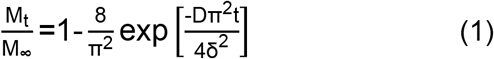

where M_t_ is the mass of released TRITC-dextran at time t, M_∞_ is the cumulative mass of TRITC-dextran, D is the effective diffusion coefficient, and 2δ is the hydrogel thickness.

The theoretical mesh size of the hydrogel compositions was calculated using the hindered solute diffusion in solvent-filled pores model as described by Equation (2).

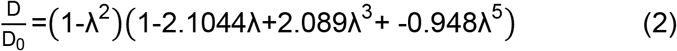

Where D_0_ is the theoretical diffusion coefficient of TRITC-dextran in free solvent and λ is a characteristic ratio of TRITC-dextran hydrodynamic diameter to theoretical average mesh size of the polymer network in the hydrogel matrix.

The hydrodynamic diameter of 4.4 kDa and 150 kDa TRITC-dextran was estimated to be 2.8 nm and 17 nm respectively. The diffusivity of 4.4 kDa and 150 kDa TRITC-dextran (D_0_) was calculated from the Stokes-Einstein equation (Equation 3) to be 1.40×10^-6^ m^2^/s and 2.09×10^-7^ m^2^/s respectively.

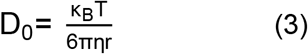

Where κ_B_ is the Boltzmann constant, T is the absolute temperature, η is the dynamic viscosity of TRITC dextran depending on its molecular weight, and r is the hydrodynamic radius of TRITC-dextran.

#### 5. Hydrogel degradation

Photocrosslinked hydrogels of varying conditions (discs of 4 mm diameter and 1 mm thickness) were swollen to equilibrium. Coomassie dye was used as a protein-specific marker. Swollen hydrogels were incubated overnight in 0.1% (w/v) Coomassie dye with gentle agitation to ensure uniform staining. Since Coomassie dye binds to proteins with high affinity, the gels were subsequently destained for 3 hours in a destaining buffer (PBS) to remove any unbound dye, ensuring that only fibrinogen-bound dye remained within the hydrogel network. PEGDA hydrogels also displayed mild staining, possibly due to diffusion-mediated physical entrapment of the dye. The destained hydrogels were transferred to multiwell plates and incubated with collagenase I (Himedia, 125 U/mg, working concentration: 100 U/mL) in PBS buffer at 37°C for total duration of 3 hours. 50 μL of the PBS buffer was collected at 15-minute intervals and replaced with equal volume of the warm collagenase I solution. The absorbance values at 595 nm was measured using a multimode plate reader (Biotek Synergy H1). PBS buffer was used as the blank control. The data obtained indicated the release of Coomassie-blue bound fibrinogen due to enzymatic degradation of the hydrogel network, as well as some diffusion-mediated release of unbound dye. The diffusion-mediated release values (based on PEGDA hydrogel condition) were subtracted from all the conditions to obtain purely degradation-mediated release values. The relative degradation for each hydrogel condition was plotted over time. The degradation rate was calculated from the average slope of the initial linear region of the relative degradation curve. A minimum of 5 hydrogel replicates per condition were measured.

#### 6. Hydrogel stiffness estimation via representative model

The complex shear moduli of the hydrogel conditions were estimated based on previously established equations related to equilibrium swelling theory and rubber-elasticity theory on polymer-solvent networks. It is described as below:

The number average molecular weight (M_n_) and degree of polymerization between crosslinks (N_c_) of the PEG-fibrinogen/PEGDA hydrogel composites were estimated based on the Equations (4) and (5).

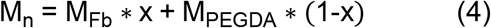

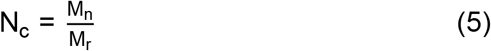

Where x = mass fraction of fibrinogen estimated in Table 1, M_Fb_ = molecular weight of bovine fibrinogen (640 kDa), M_PEGDA_ = molecular weight of PEGDA (10 kDa), M_r_ = molecular weight of the repeating polymeric unit.

The initial polymer volume fraction (Φ_o_) of the hydrogel formulations were estimated using known polymer and water densities using Equation (6).

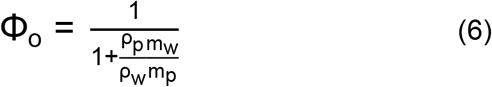

Where ρ_p_ = density of polymer (1.18-1.35 g/cm^3^), ρ_w_ = density of water (1.00 g/cm^3^), m_p_ = polymer mass and m_w_ = mass of water.

The relaxed polymer volume fraction (Φ_r_), swollen polymer volume fraction (Φ_s_), and reference ratio (θ) were calculated based on Φ_o_ and N_c_ values according to Equations (7-9).

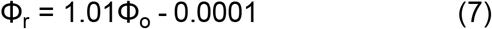

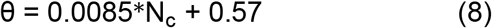

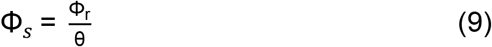

The effective molecular weight between crosslinks 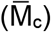 for each hydrogel formulation was calculated using Equation (10).

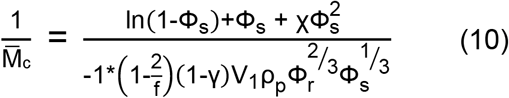

Where f = effective junction functionality (assumed to be ∞), γ = frequency of chain-end defects (assumed to be 0), V1 = Molar volume of solvent, water (18 g/mol), χ = Flory’s polymer-solvent interaction parameter (0.46-0.50).

The complex shear modulus (G*) was calculated from the rubber elasticity theory using Equation (11)

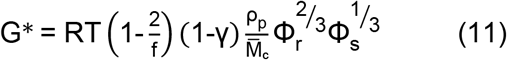

